# Aromatic residues in mobile regions distal to the active site support the closed conformation of *E. coli* DXPS

**DOI:** 10.1101/2025.10.24.684457

**Authors:** Lydia J. Kramer, Steven L. Austin, Ananya Majumdar, Noah D. Smith, H. Lee Woodcock, Caren L. Freel Meyers

## Abstract

The essential bacterial enzyme 1-deoxy-D-xylulose 5-phosphate synthase (DXPS) is absent in humans, making the enzyme an attractive antimicrobial target. Its product DXP sits at a metabolic branchpoint between the biosynthesis of pyridoxal phosphate (PLP), thiamin diphosphate (ThDP), and isoprenoids. DXP is formed via decarboxylation of pyruvate and subsequent carboligation with D-glyceraldehyde-3-phosphate (D-GAP) in a ThDP-dependent manner. In the current mechanistic model, DXPS follows a ligand-gated mechanism. Pyruvate reacts with ThDP to form C2⍺-lactylThDP (LThDP) which coincides with a shift to a closed conformation. The flexible “spoon” and “fork” motifs become ordered, situating the catalytic residue H299 within the active site which supports LThDP persistence and the closed conformation of the E-LThDP complex until binding of D-GAP. Our goal is to understand the molecular basis for stabilization of the E-LThDP complex in its closed conformation in the absence of D-GAP. We propose the conserved aromatic residues Y288, F298, and F304 in the *E. coli* DXPS spoon and fork motifs form a cluster upon transition from the open to closed form to position H299 within the active site, necessary for LThDP persistence. Here, we conducted mutagenesis studies to elucidate the roles of Y288, F298, and F304 in conformational cycling and catalysis. On each variant, the conformational equilibrium favored an open state, hindered intermediate formation and persistence, and promoted intermediate release from the active site. Our results support a model in which conserved aromatic residues within the mobile, sequence-diverse spoon and fork motifs promote the closed conformation and support catalysis.

## Introduction

The thiamin diphosphate (ThDP)-dependent enzyme 1-deoxy-D-xylulose 5-phosphate synthase (DXPS) catalyzes the decarboxylative condensation of pyruvate and D-glyceraldehyde-3-phosphate (D-GAP) to form DXP (Figure 1).^1–3^ The product DXP is a branchpoint metabolite required for the synthesis of essential isoprenoids and cofactors (ThDP and pyridoxal phosphate, PLP).^1,2,4,5^ We showed that DXPS functions in bacterial adaptations that rely upon isoprenoids or vitamins, demonstrating a role of DXPS in the PLP-dependent detoxification of the bacteriostatic host metabolite D-serine by uropathogenic *E. coli* (*Ec*) in the urinary tract.^6^ While the DXPS-catalyzed reaction is reminiscent of reactions catalyzed by other ThDP-dependent enzymes, including the E1 subunit of pyruvate dehydrogenase (PDH) and transketolase (TK), DXPS has a distinct homodimer domain arrangement and ligand-gated mechanism.^7–9^ DXPS also exhibits relaxed substrate specificity, catalytic promiscuity, a large active site volume, and conformational diversity, all of which are characteristics of multifunctionality.^10,11^ Together with its ligand-gated mechanism, these features may facilitate DXPS activities other than the synthesis of DXP, which may aid in pathogen adaptation.^12–19^ Selective inhibition of DXPS therefore has the potential to hinder microbial adaptation during infection. The importance of DXPS in bacterial metabolism and pathogen adaptation, as well as its absence in humans, makes it an attractive anti-infective target.^12,13,20–36^

**Figure 1:**
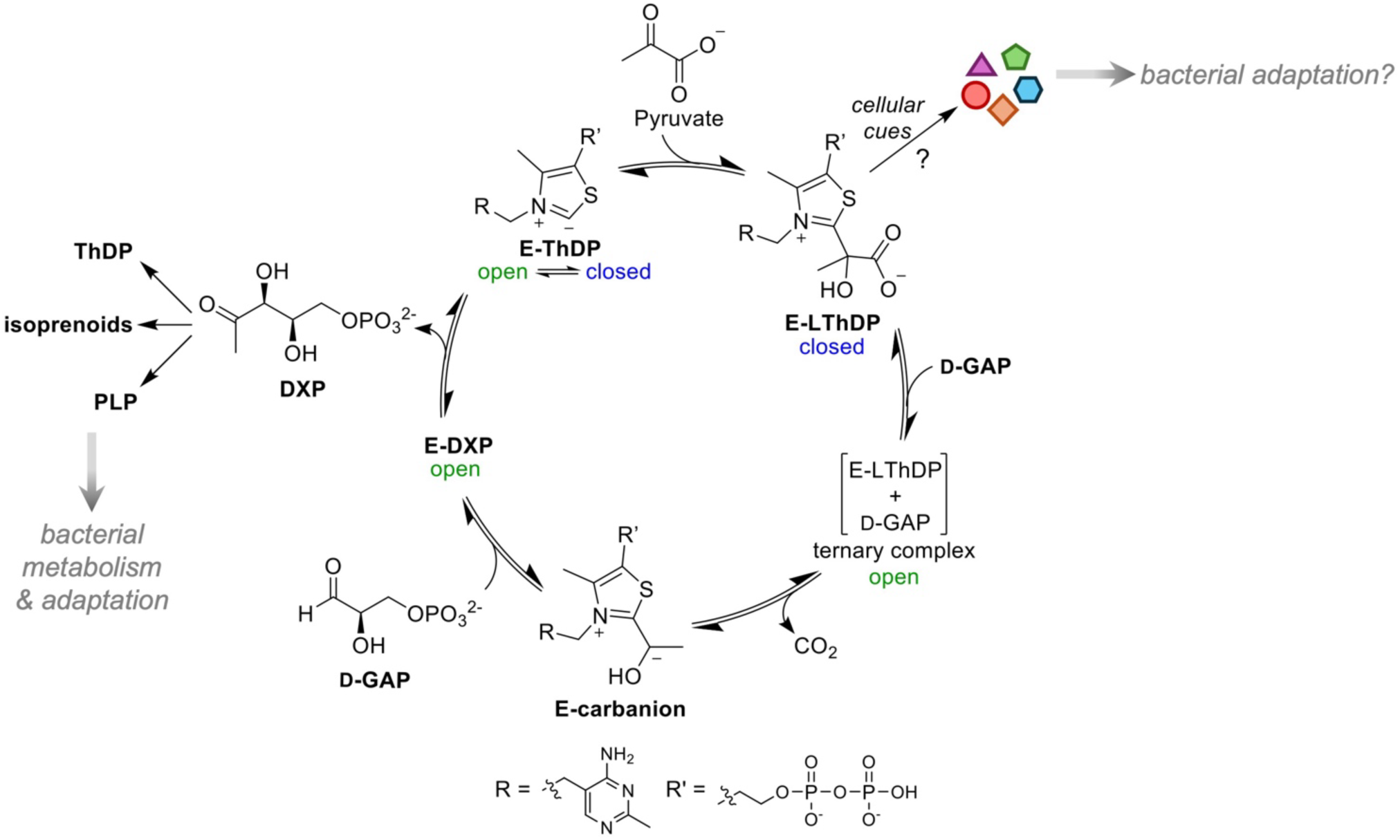
DXPS-catalyzed synthesis of the branchpoint metabolite DXP. DXPS catalyzes the ThDP-dependent decarboxylation of pyruvate followed by carboligation with D-GAP to produce the branchpoint metabolite DXP via a ligand-gated mechanism. Catalysis is closely linked to conformational changes. The E-LThDP complex is predicted to exist in a closed conformation with LThDP coordinated by an active site network. D-GAP binding induces LThDP decarboxylation coinciding with a shift to the open state. Canonical and alternative functions of DXPS may aid in bacterial adaptation.

Unique among ThDP-dependent enzymes, the ligand-gated mechanism of DXPS has been exploited for selective inhibitor design.^23–25,37–39^ Upon DXPS binding to its donor substrate pyruvate, the pre-decarboxylation intermediate C2α-lactylThDP (LThDP) forms (E-LThDP, Figure 1). LThDP persists on DXPS in the absence of the co-substrate D-GAP, stabilized by an active site network consisting of *Ec*H431-D427-R99-H49-H299 (Figure 2, left).^14,40^ Upon binding of D-GAP, LThDP is activated for decarboxylation, forming a carbanion (E-carbanion, Figure 1) that subsequently reacts with D-GAP to form DXP.^41^ Thus, D-GAP serves two roles in this DXP-forming mechanism, (1) as a “trigger” of LThDP decarboxylation in the gated step and (2) as an acceptor in the following carboligation event. Our lab has identified alternate donors, acceptors, and triggers of LThDP decarboxylation, which when taken together with the ligand-gated mechanism, suggest that DXPS can sense and respond to changes in the bacterial metabolic environment.^12,14,17–19,32,42^ It is also plausible that changes in substrate availability may be sensed by DXPS, facilitating alternative product formation that could aid in pathogen adaption. Thus, DXPS may play important roles in host-pathogen interactions during infection that could be selectively targeted, underscoring the importance of understanding its mechanism.

**Figure 2.**
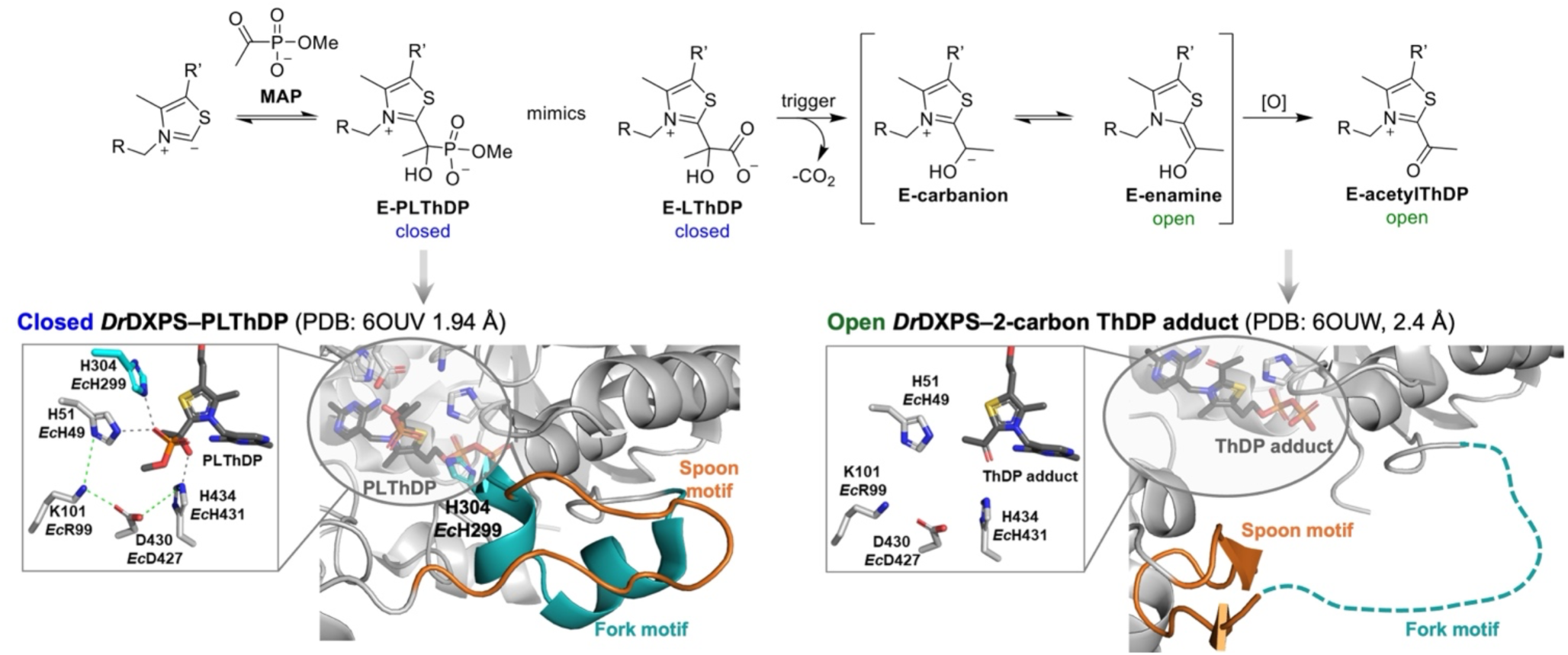
DXPS conformational changes associated with LThDP formation and decarboxylation. **Left:** The PLThDP adduct formed upon addition of MAP (E-PLThDP, top left) mimics the natural LThDP intermediate (E-LThDP, top center). PLThDP formation coincides with the closed conformation and an intact active site network (lower left, PDB: 6OUV). In the closed conformation, the ordered spoon (orange) and fork (cyan) motifs fill most of the active site cleft. **Right:** The post-decarboxylation structure (lower right, PDB: 6OUW) shows an open conformation with the spoon motif positioned to one side of the active site cleft, and a disordered fork motif. This open conformation shows a planar 2-carbon ThDP adduct bound to *Dr*DXPS, which could be either the E-enamine or E-acetylThDP (E-acetylThDP shown).

Progression along the DXPS reaction coordinate is driven by dramatic conformational changes. In the absence of substrates, DXPS exists in a conformational equilibrium between open and closed states, with the closed state favored.^9,13,15,43^ The DXPS closed confirmation was observed through incubation with the pyruvate analog methyl acetylphophonate (MAP), which forms a stable enzyme-bound phosphonolactyl ThDP adduct (E-PLThDP) that mimics the natural LThDP intermediate (Figure 2, left). Formation of E-PLThDP coincides with a shift in the conformational equilibrium to the closed form, detected by X-ray crystallography and hydrogen-deuterium exchange mass spectrometry (HDX-MS) where protection from deuterium exchange in regions near the active site by is observed.^13,15,33^ A shift to the open conformation is detected upon addition of D-GAP to the E-PLThDP complex derived from MAP, and in the post-decarboxylation state (Figure 2).^33^ Crystallography studies of *D. radiodurans* (*Dr*) DXPS support that LThDP formation coincides with a shift in the conformational equilibrium in which a flexible loop containing two motifs - termed the fork and spoon motifs - become ordered, filling the active site cleft in a ‘closed’ form (Figures 2, left, and S1a).^13,33^ This structural shift positions key catalytic residue H299 in close proximity to LThDP and completes assembly of the active site network (*Ec*H431-D427-R99-H49-H299, Figure 1 top, right inset) which prevents LThDP activation on DXPS in the absence of D-GAP or other small molecule triggers of LThDP activation.^14,19,43^ We have proposed that binding of a trigger molecule disrupts the active site network, promotes disorder of the fork motif, and causes repositioning of the spoon motif to one side of the active site, producing an open state. This conformational shift coincides with removal of H299 from the active site and LThDP activation (Figures 1 bottom inset, 2 (right), and S1b).^13,33^ Based on these insights, we suggest that the reactivity of active site intermediates is dictated by movements in the fork/spoon motifs. However, it remains unknown how the flexible spoon and fork motifs promote the closed conformation to support a stable E-LThDP complex in the absence of a trigger molecule.

Here, we have identified a conserved cluster of aromatic residues in the spoon/fork motifs (Y288, F298, and F304 on *Ec*DXPS) that supports catalysis and plays a role in the persistence of LThDP on DXPS, likely by ensuring correct positioning of the catalytic H299 residue within the active site. We hypothesize that the aromatic cluster supports the closed conformation in the absence of a small molecule trigger, maintaining an intact active site network which promotes LThDP persistence. We demonstrate that substitution of any of these aromatic residues reduces the catalytic efficiency of DXPS by hindering LThDP formation. Disruption of the aromatic cluster also shifts DXPS to an open state, destabilizes LThDP, and promotes release of the post-decarboxylation intermediate from the active site. Despite the location of the aromatic cluster outside of the active site, coordination of these residues appears to be required for catalysis by positioning H299 within the active site. While the spoon and fork motifs share low overall sequence conservation between DXPS homologs, it is possible there is a conserved role of aromatic clustering on DXPS homologs, with differences in cluster composition possibly pointing to species differences in ligand gating mechanisms. Understanding the roles of flexible loops on DXPS catalysis may inform studies of alternative enzyme functions in bacteria and will guide the development species-specific DXPS inhibitors.

## Materials and Methods

### General Methods

All reagents were purchased from commercial sources unless otherwise noted. All primers for mutagenesis were purchased from Integrated DNA Technologies. Chemicals used in the described experiments were obtained from Millipore Sigma unless otherwise noted. 4-(2-Hydroxyethyl)-1-piperazineethanesulfonic acid (HEPES) and LB broth were purchased from Fisher Scientific. *E. coli* DXPS and IspC recombinant enzymes were overexpressed and purified as reported previously.^14,15,17,18,32,37,39,43,45^ A vinyl Coy Laboratory Products anaerobic chamber (Grass Lake, MI, USA) was used for anaerobic experiments. A BioTek Epoch 2 microplate reader was used for spectrophotometric analysis in kinetic experiments. Steady-state and stopped-flow circular dichroism (CD) experiments were performed using an Applied Photophysics Chirascan V100 CD spectrometer (Surrey, U.K). For pre-steady state experiments, the CD spectrometer was fitted with a Chirascan SF3 accessory (Surrey, U.K.). Anaerobic solutions for CD experiments conducted outside of the anaerobic chamber were prepared within the chamber and transferred to airtight cuvettes or vials fitted with septa for the duration of the experiment. SDS-PAGE gels were imaged using a Li-COR Odyssey CLx imager. NMR spectra of the synthesized HEThDP standard were acquired on a JEOL JNM-ECZL500R spectrometer. A Bruker Avance spectrometer outfitted with a TCI cryogenic probe containing a cryo-cooled ^13^C preamplifier was used to detect enzymatically-generated HEThDP by dimensional ^13^C and ^1^H NMR.

### Molecular Dynamics Simulations of *Dr*DXPS-PLThDP

The PLThDP-bound *Dr*DXPS crystal structure (PDB: 6OUV) was selected to provide the initial coordinates for molecular dynamics simulations.^33^ A region of DXPS (*Dr*DXPS 200−243) is unresolved in one monomer and only partially resolved in other crystal structures. To obtain a complete structure of E-PLThDP, we used the full-length FASTA sequence as input to AlphaFold v2 (AF) to subsequently obtain coordinates for the unresolved region.^46^ To retain the known crystal coordinates, the AF structure was first aligned to the crystal structure, and the missing region was then grafted onto the crystal structure to produce the complete *Dr*DXPS structure for simulation. The CHARMM-GUI web server was utilized to protonate histidine residues and solvate the system in a cube with side lengths of 120 Å containing 47662 TIP3 water molecules with 168 potassium and 135 chloride ions for neutralization.^47^ All molecular dynamics calculations were performed using CHARMM.^48^ Parameters for PLThDP were generated with CGenFF.^49^ The CHARMM36 force field including CMAP corrections was applied to protein and ions. Nonbonded interactions were truncated at 16 Å. Electrostatic interactions were treated with the particle-mesh Ewald method. Equilibration was performed by applying harmonic restraints with 1.0 and 0.1 kcal/mol force constants applied to the backbone and side chains, respectively. 50 steps of steepest descent minimization followed by 50 steps of adopted basis Newton−Raphson minimization were performed before 0.125 ns in the NVT ensemble with a time step of 1 fs at 303.15 K. A production run of 250 ns was performed with a 2 fs time step in the NPT ensemble, utilizing the Nose-Hoover thermostat and Langevin Piston algorithm to maintain the temperature and pressure (1 atm) respectively. The SHAKE algorithm was applied to constrain all bonds to hydrogen. Coordinates were saved every 4 ps. This production run was performed to ensure structural stability in preparation for simulation with Anton 2, where we collected a 23 μs trajectory in the NPT ensemble at 310k.^50^ All analyses were performed with in-house scripts to carry out principal component analysis, free energy surface construction, and various other trajectory measurements.

### Generation of DXPS variants

Using a method adapted from the New England Biolabs Q5 Site-Directed Mutagenesis protocol, we generated F298A/L, Y288A/F/Q, and F304A/L *Ec dxs*-pET37b constructs. The *dxs*-pET37b plasmid, purified from *E. coli* Top 10 competent cells using a Qiagen QIAprep Spin Miniprep kit, was used as the template. Polymerase chain reaction (PCR) mixtures (25 µL volume) contained 1X Q5 Hot Start High-Fidelity Master Mix (NEB), 1 ng/µL *dxs*-pET37b template plasmid, and 10 µM forward and reverse primers (Table S1) designed using the NEBaseChanger primer design tool and purchased from IDT. The PCR was conducted as follows: 30 s at 98 °C; 25 cycles of 10 s at 98 °C, 30 s at the annealing temperature specific to each primer set (Table S1), and 10 min at 72 °C for elongation; followed by a final elongation for 2 min at 72 °C. PCR products were stored at 4 °C until use in the (10 µL volume) Kinase-Ligase-Digestion (KLD) reaction (NEB) which were made from 1 µL PCR product, 1X KLD reaction buffer, 1X KLD enzyme mix, and incubated for 5 min. The KLD product was transformed into *E. coli* Top 10 competent using the established heat shock protocol (5µL reaction product and 50µL competent cells) and positive transformants were selected with kanamycin (50 µg/mL).^31,43^ Constructs were purified from colonies and sequenced by Genewiz to confirm the desired substitution and presence of the C-His_8_ tag.

Plasmids were then transformed into competent *E. coli* BL21 (DE3) cells for protein overexpression and subsequent purification. Briefly, 2 L cultures containing BL21 (DE3) cells harboring *Ec* WT and variant DXPS constructs were grown with shaking at 37 °C for 2 h, at which point the temperature was reduced to 25 °C. When cultures reached OD_600_ of 0.6 (log phase of bacterial growth), IPTG was added to a final concentration of 100 µM to induce *dxs* overexpression. Cultures were grown with shaking for ∼20 h before centrifugation at 18,000 x g for 12 min at 4 °C. The supernatant was removed; the cell pellets were flash frozen in liquid nitrogen and stored at −80 °C until protein purification. The cell pellet was resuspended in 100 mL lysis buffer (400 mM NaCl; 50 mM 100 mM 2-amino-2-(hydroxymethyl)1,3-propanedio (Tris), pH 8; 10% glycerol; 1 mM phenylmethylsulfonyl fluoride (PMSF) containing 1 mM Tris(2-carboxyethyl)phosphine (TCEP), 1 mM ThDP, 1X protease inhibitor cocktail (PIC), 20 mM MgCl_2_, and 6 µL DNase/40 mL). The resuspended cultures were lysed by passage through the Microfluidizer^TM^ LM10 four times. The lysate was centrifuged at 39,000 x g for 30 min at 4 °C and ∼100 mL supernatant was combined with 7 mL Ni-NTA resin, previously washed with lysis buffer. Imidazole was added to a final concentration of 5 mM to reduce nonspecific protein binding to the resin. This suspension was placed in a 4 °C cold room and gently rocked for 1 h to promote DXPS binding to the Ni-NTA resin. The DXPS-bound Ni-NTA resin was then centrifuged at 4,000 x g for 10 min at 4 °C to separate the resin from the supernatant containing undesired/unbound protein. The DXPS-bound resin was loaded into a gravity column and washed with 8 mL each of 5, 20, 40, and 60 mM imidazole (prepared in lysis buffer) to remove nonspecifically-bound protein from the resin. DXPS was eluted in a stepwise imidazole gradient (4 mL each of 100 mM, 250 mM, and 4 x 4 mL of 500 mM imidazole solutions prepared in lysis buffer). Fractions obtained from 250 and the first three 500 mM imidazole elutions were pooled (16 mL final volume) and diluted 8-fold (128 mL final volume) into anion exchange buffer A (50 mM Tris, pH 8, and 10% glycerol). Precipitate was removed by centrifugation at 39,000 x g for 30 min at 4 °C. The supernatant was loaded at 2 mL/min onto Cytiva HiTrap HP Q columns connected in series (2 x 5 mL columns). Two column volumes (20 mL) of anion exchange buffer A containing 1 mM TCEP were passed through the columns to remove nonspecifically-bound proteins. DXPS was then eluted using a gradient of 0-100% anion exchange buffer B (50 mM Tris, pH 8; 10% glycerol; 1 M NaCl; and 1 mM TCEP) in buffer A, over 10 column volumes using a 2.5 mL/min flow rate (100 mL over 40 min). The eluted protein was pooled and dialyzed overnight in 1 L dialysis buffer (50 mM Tris, pH 8; 10 mM MgCl_2_; 100 mM NaCl, 10% glycerol, 1 mM ThDP) at 4 °C. The following day protein was dialyzed at 4 °C for 4h in 1 L of fresh dialysis buffer. Protein concentration was determined by Bradford assay, and the protein stock was flash frozen in liquid nitrogen and stored at −80 °C.

### Determination of DXPS variant secondary structure

The secondary structures of WT and variant DXPS enzymes were determined using CD, as reported previously.^15,31,43^ Each enzyme was buffer exchanged (25 mM Tris, pH 8, 10 µM ThDP, 1 mM MgCl_2_, and 100mM NaCl) prior to far-UV CD measurements. The absorbance at 280 nm of a denatured sample, consisting of 400 µL of guanidine-HCl (7.5 M stock) with 100 µL of enzyme (stocks ranging from 91-168 μM), was measured, and the concentration of DXPS monomers was determined from absorbance using calculated extinction coefficients (ε) from equation 1.

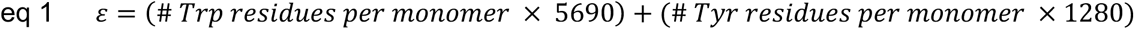

The predicted extinction coefficient for WT, F298L, and F304L (47080 M^−1^cm^−1^) was used as published previously.^15,43^ However, the Y288Q concentration was calculated using a predicted 45800 M^−1^cm^−1^ extinction coefficient due to the substitution of one of nineteen Tyr residues per monomer. Secondary structures were analyzed in triplicate using 3 µM non-denatured WT or variant DXPS enzyme in exchange buffer at 25 °C in a 5 mm cuvette, by measuring the CD signal over the range 250-180nm, with 1 nm step, and 2 s averaging time.

### Determination of apparent T_m_ of DXPS variants

The T_m_ of each DXPS variant was determined by monitoring the secondary structure via CD by thermal denaturing, as described previously.^15,31,43^ Each enzyme sample was buffer exchanged, and the concentration was determined as described above for “determination of DXPS variant secondary structure”. To determine the apparent melting temperature, the ⍺-helical content of DXPS was monitored by the CD signal at 222 nm from 20–88 °C, at a rate of 1 °C/min, a 0.2 °C tolerance, a 20 s settling time at each temperature, and a 20 s averaging time at each temperature with stirring. The CD signal at 222 nm was plotted versus temperature, and the apparent T_m_^app^ was determined as the maximum of the first derivative of this plot. T_m_^app^ determinations were conducted in triplicate.

### DXPS-IspC coupled assay

DXPS-dependent formation of DXP was detected by the DXPS-IspC coupled assay and used for kinetic characterizations (below). The general method used for the DXPS-IspC coupled assay is as follows: Enzyme reactions were performed in a 96-well plate. Enzyme solution (180 µL) containing DXPS (concentrations specified below), IspC (2 µM final concentration), and nicotinamide adenine dinucleotide phosphate (NADPH, 200 µM) in reaction buffer (100 mM HEPES pH 8, 2mM MgCl_2_, 5mM NaCl, and 1mM ThDP) was incubated at 30 °C before initiation of the reaction by adding 20 µL of a 10x substrate solution containing both pyruvate and D-GAP (200 µL final volume). The depletion of NADPH was monitored at 340 nm at 30 °C using a BioTek Epoch 2 microplate reader. The initial velocity of NADPH disappearance was used to calculate the initial velocity of DXP formation.

### Kinetic characterization of DXP formation on DXPS variants

Kinetic parameters (*k*_cat_, *K*_m_, and *k*_cat_/*K*_m_) were determined using DXPS-IspC-coupled assay as described above in “DXPS-IspC coupled assay”. The final DXPS enzyme concentration was 100 nM (WT and Y288F) or 1.5 µM (Y288A/Q, F298A/L, and F304A/L). The *K*_m_ values for pyruvate and D-GAP were determined individually by varying one substrate while holding the other constant at a saturating concentration. Pyruvate concentrations were varied over the range 0-500 µM (WT, Y288F, F298A/L, and F304A/L) or 0–10 mM (Y288A/Q) while maintaining D-GAP concentration at 500 µM. When D-GAP was the varied substrate (0–500 µM for WT and all DXPS variants), pyruvate concentration was maintained at 500 µM (for WT, Y288F, F298A/L, and F304A/L) or 5 mM (for Y288A/Q). Reactions were monitored at 340 nm for 10 minutes, measuring the absorbance every 20 seconds. The initial velocity of DXP formation was determined from the linear portion of the timecourse and plotted versus substrate concentration in GraphPad Prism version 10 and fit to equation 2, where v_i_ is the initial velocity, V_max_ is the maximum velocity, [S] is the substrate concentration and *K*_m_ is the concentration of substrate required to reach half maximum velocity.

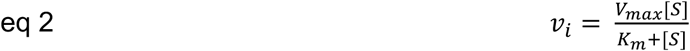

Turnover efficiency (*k*_cat_/*K*_m_) was determined by measuring the slope of the initial, linear region of the Michaelis-Menten curve (v/[E] versus [S]). Two technical replicates were conducted in each experiment and were repeated for a total of four replicates. Kinetic parameters are reported as the mean with standard error determined from four replicates for *K*_m_ and *K*_m_/*k*_cat_ or 8 replicates for *k*_cat_.

### CD analysis of aminopyrimidine (AP) and 1’,4’-iminopyrimidine (IP) tautomers of ThDP on DXPS variants

Since LThDP is readily activated in the presence of O_2_, all experiments were performed anaerobically, as reported previously.^15,43^ WT or variant DXPS (30 µM) samples were prepared anaerobically at 4 °C in buffer (50 mM HEPES, pH 8, 100 mM NaCl, 1 mM MgCl_2_, 0.2 mM ThDP) to promote the AP tautomer. The buffer was prepared using anaerobic stocks (prepared from solids and deoxygenated water). Anaerobic conditions were established and maintained through sample removal, with samples transferred in airtight, septum-capped vials or 10 mm path-length cuvettes (Starna cells, 1-Q-10-ST-S). To assess differences in the LThDP IP signals between WT and variant DXPS, anaerobic pyruvate solutions (150 mM or 692 mM stocks) were added to enzyme using a gas-tight Hamilton syringe for final pyruvate concentrations of 5x*K*_m_^Pyruvate^ (200 µM for WT DXPS, 5 mM for Y288Q and the dialysis buffer control, 300 µM for F298L, 200 µM for F304L), and the CD signal was recorded over the range 290–400 nm. All CD spectra were collected on 1.5 mL samples in a 10 mm path-length cuvette. CD scans were recorded at 25 °C from 290 nm to 400 nm with a 1 nm step and 0.5 s averaging time. Experiments were conducted in duplicate.

### Detecting accumulation of the PLThDP adduct derived from MAP on WT DXPS and variants

*Time-resolved CD to measure PLThDP formation:* Previously acetylphosphonate-based inhibitors of DXPS have been examined using CD by measuring the CD signal corresponding to the PLThDP adduct at 294 nm.^39^ This CD signal is typically broad; thus, we chose to monitor the CD_313_ signal used to monitor LThDP formation, given the pre-steady state limitation that lower wavelengths result in high voltages that influence the measured CD signal. To monitor PLThDP accumulation, 60 µM WT or variant DXPS was prepared in buffer (50 mM HEPES, pH 8, 100 mM NaCl, 2 mM MgCl_2_, and 2 mM ThDP). Separately, a 10 mM MAP solution was prepared in the same buffer. The CD was fitted with the stopped-flow SF3 accessory to enable rapid mixing of enzyme and MAP solutions (30 µM enzyme and 5 mM MAP final concentrations), and the change in the CD_313_ signal was measured over 50 s at 10 °C. Data from 7 repetitive shots were averaged and plotted.

*PLThDP formation over a long timecourse:* Using CD, the formation of PLThDP was monitored over a longer timescale in order to determine maximal signal amplitudes correlating to PLThDP on WT and DXPS variants. WT or variant DXPS (30 µM) was added to buffer (50 mM HEPES, pH 8, 100 mM NaCl, 1 mM MgCl_2_, 0.2 mM ThDP). Experiments were initiated by the addition of 5 mM MAP, and the CD_313_ signal was measured every 10 s with 0.5 s averaging time, over 13 min. Since DXPS variants display unique AP signal shifts and amplitudes relative to WT DXPS, the change in the PLThDP CD_313_ signal was normalized to the average AP signal at 313 nm (calculated from two back-to-back scans) in each case (PLThDP CD_313_ signal – AP CD_313_ signal) and plotted as a function of time. Experiments were performed in duplicate.

### Limited trypsinolysis

WT and variant DXPS (9 µM) were incubated on ice in 100 mM HEPES, pH 8, 100 mM NaCl, 2 mM MgCl_2_, 1 mM ThDP, and 1% glycerol for 45 min in the presence or absence of 5 mM MAP. Digestion was initiated by the addition of 4.5 ng/µL trypsin, and the reaction mixture was incubated on ice for the duration of the experiment (60 min). Aliquots (10 µL) were quenched into cold 2x sodium dodecyl sulfate (SDS) loading dye (100mM Tris, pH 6.8; 4% (w/v) SDS; 0.2% (w/v) bromophenol blue; 20% (w/v) glycerol; and 200 mM dithiothreitol) at 1, 5, 15, 30, 45, 60 min, vortexed, and flash frozen in liquid nitrogen prior to SDS-PAGE analysis. SDS-PAGE gels were stained with Colloidal Coomassie G-250 stain and imaged using a LI-COR Odyssey CLx. Full-length DXPS (68 kDa) and digest products were quantified using Fiji. In this analysis, bands near 44 kDa were combined for analysis and bands near 34 kDa were combined for analysis; these additional bands near 44 and 34 kDa likely arise from the cleavage of nearby Lys residues. The percent band density for each group of peptide products was determined by equation 3.

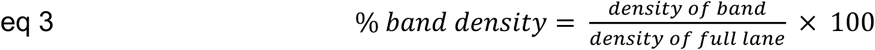

Experiments were conducted in triplicate, and the standard errors of the percent band densities were calculated. A Welch t test was used to determine statistical significance of the difference between ∼44 and ∼34 kDa (including the nearby bands observed in the DXPS variant gels).

### Observing LThDP persistence by CD

LThDP persistence on WT and variant DXPS was assessed by monitoring the stability of the IP CD_313_ signal corresponding to LThDP. All solutions were degassed in the anaerobic chamber at 4 °C before preparation of a 30 µM enzyme solution in buffer (50 mM HEPES, pH 8, 100 mM NaCl, 1 mM MgCl_2_, 0.2 mM ThDP), and samples were transferred out of the chamber in air-tight, septa-capped cuvettes. An initial scan was recorded at 25 °C from 290-400 nm, using a 1 nm step and 0.5 s averaging time, to detect the AP signal. LThDP formation was initiated by addition of pyruvate at 5x*K*_m_^Pyruvate^ (200 µM for WT DXPS, 5 mM for Y288Q, 300 µM for F298L, 200 µM for F304L) from an anaerobic pyruvate solution using a gas-tight Hamilton syringe, and the solution was mixed by inverting the cuvette. The persistence of the CD_313_ LThDP signal and its depletion were monitored at 25 °C until the depleting signal reached equilibrium (50 or 70 min), with scans (290-400 nm) collected every 5 min, using a 1 nm step, and 0.5 s averaging time. IP signal depletion was evaluated by calculating the percent change in CD_313_ in equation 4.

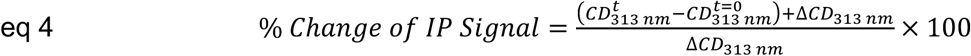

Experiments were conducted in duplicate for each variant and WT DXPS.

### NMR detection of bicarbonate and 2-(1-hydroxyethyl)ThDP (HEThDP) upon disruption of the aromatic cluster

NMR experiments were performed as previously published with minor modifications.^19,32,43^ DXPS-catalyzed reactions were performed and quenched in the anaerobic chamber to prevent LThDP decarboxylation induced by molecular oxygen. Reaction mixtures were prepared from degassed enzyme stocks, and buffer and salt solutions were prepared from solid stocks suspended in degassed water. WT or variant DXPS (5 µM final concentration) was added to buffer (50 mM HEPES, pH 8, 100 mM NaCl, 2 mM MgCl_2_, 1 mM ThDP, and 5 % D_2_O). LThDP formation was initiated by adding 7 µL of 50 mM [^13^C_3_]pyruvate to give a final [pyruvate] of 500 µM and a final reaction volume of 700 µL. Samples were incubated at ambient chamber temperature (∼28 °C) for 20 min prior to quenching by boiling at 95 °C for 5 min. Solutions were transferred to 3 kDa microfuge filter tubes, removed from the chamber, and centrifuged at 14,000 x g for 10 min to remove DXPS. To 600 µL of filtrate, 15 µL of 10 mM gadobutrol (MedChem Express, NJ) was added to enhance the bicarb signal.^50^ Samples were stored at 4 °C until NMR spectra acquisition which was performed the same day. Spectra were acquired at 150 MHz (^13^C), at 25 °C, using a 35° ^13^C excitation pulse, 1,320 scans/FID, 150 ms acquisition time/FID, and a 3s relaxation between scans.

### Synthesis of HEThDP standard

A HEThDP standard was prepared using the procedure of Gruys, Halkides, and Frey,^51^ with minor modifications. To a solution of ThDP (692 mg, 1.0 equivalent, 1.5 mmol) in 5 mL distilled water was added freshly distilled acetaldehyde (2 mL, 24 equivalents, 36 mmol). While stirring the mixture cooled in an ice water bath to minimize boiling of acetaldehyde, the pH was adjusted to 8 using 5 M NaOH and then increased to pH 8.5–9 using 1 M NaOH. The reaction flask containing the pale-yellow solution was sealed with a septum, and the solution was stirred at 45 °C for 2 hours. The reaction was then cooled over ice, and the pH was increased to 8.5–9 by dropwise addition of 1 M NaOH. The flask was resealed, and the solution was stirred at 45 °C for 1 hour. After 3 hours, the ratio of HEThDP/ThDP was 3:1 as determined by ^1^H-NMR.

The reaction was quenched by dropwise addition of conc. HCl to pH 8. The reaction mixture was concentrated by rotary evaporation at 50 °C to remove water and acetaldehyde. The remaining 1 mL of bright yellow solution was loaded onto a Sfär C18 column that was equilibrated with 1% aqueous formic acid, and the mixture was eluted with a stepwise gradient of formic acid (A) and acetonitrile (B): 1) 100% A, 4 column volumes (CV); 2) 0-30% B over 5 CV; 3) 30-100% B over 2 CV; 4) 100% B for 3 CV. Fractions containing HEThDP, as determined by ^1^H NMR, were pooled and concentrated by rotary evaporation to a final volume of 1 mL. The solution was diluted with 3 mL of 0.1 M HCl and lyophilized to afford 496 mg of a white salt that contained 3:1 HEThDP/ThDP. Trace formic acid was removed by dissolving the salt in water followed by sublimation *in vacuo*. This material was used as a standard in NMR experiments (below), without further purification.

### HEThDP product identification by ^1^H NMR

Variant DXPS samples were prepared as described above (“NMR detection of bicarbonate and HEThDP upon disruption of the aromatic cluster”) and stored overnight at 4 °C prior to NMR spectra acquisition. A solution of HEThDP (15 mg/mL) was prepared and used as a standard in spike experiments in which 7 µL of this solution was added to the Y288Q sample, 4 µL was added to the F298L and H299A samples, and 8 µL was added to the F304L sample. ^1^H NMR spectra were acquired before and after addition of the HEThDP standard solution to confirm the identity of the ThDP adduct released from the active sites of DXPS variants.

## Results

### Sequence and structural alignments reveal conserved aromatic residues in the spoon and fork motifs

Two mechanistically relevant conformations have been identified on *Dr*DXPS using X-ray crystallography and HDX-MS studies in which the most dramatic structural changes were found to occur within the spoon and fork motifs. In the open conformation, the spoon motif is positioned to one side of the active site and the fork motif is disordered. In contrast, in the closed conformation, the spoon and fork motifs are positioned over the active site with H299 poised to complete the active site network containing *Ec*H431-D427-R99-H49-H299 (Figures 2 (left), and S1a).^13,15,33,43^ This network functions to stabilize E-LThDP, preventing decarboxylation by delocalizing charge from the LThDP carboxylate and minimizing strain imposed on the intermediate.^43^ How the highly mobile, sequence-variable spoon and fork regions organize in the closed conformation to support LThDP persistence is not understood. Toward addressing this question, we conducted a sequence alignment of sixteen bacterial DXPS homologs in these regions using Clustal Omega (Figure 3a),^52^ and referenced to the closed *Dr*DXPS structure (PDB: 6OUV). Interestingly, we identified an aromatic cluster within these mobile regions that is conserved on many DXPS homologs. While three aromatic residues appear to be most common, some species vary in aromatic residue number within this cluster. For example, *Ec*DXPS contains three aromatic residues (Y288, F298, and F304), while *Dr* and *Pa*DXPS contain five and two aromatic residues, respectively (Figure 3a). *Ec*Y288 was the most highly conserved residue of the three with all sixteen homologs containing either Tyr or Phe at this sequence/structural position. In homologs with fewer than three aromatic residues, isoleucine or leucine were common replacements. We predicted aromatic residues in this mobile, solvent-exposed region function to maintain the closed conformation of E-LThDP until binding of the trigger molecule. Examination of the *Dr* PLThDP-bound DXPS crystal structure in the closed conformation (PDB: 6OUV) revealed these spoon and fork aromatic residues are in close proximity, with center-to-center distances ranging from 3.9 to 9.5 Å (Figure 3b).^33^ Thus, hydrophobic and π-stacking interactions between these mobile motifs are possible.

**Figure 3:**
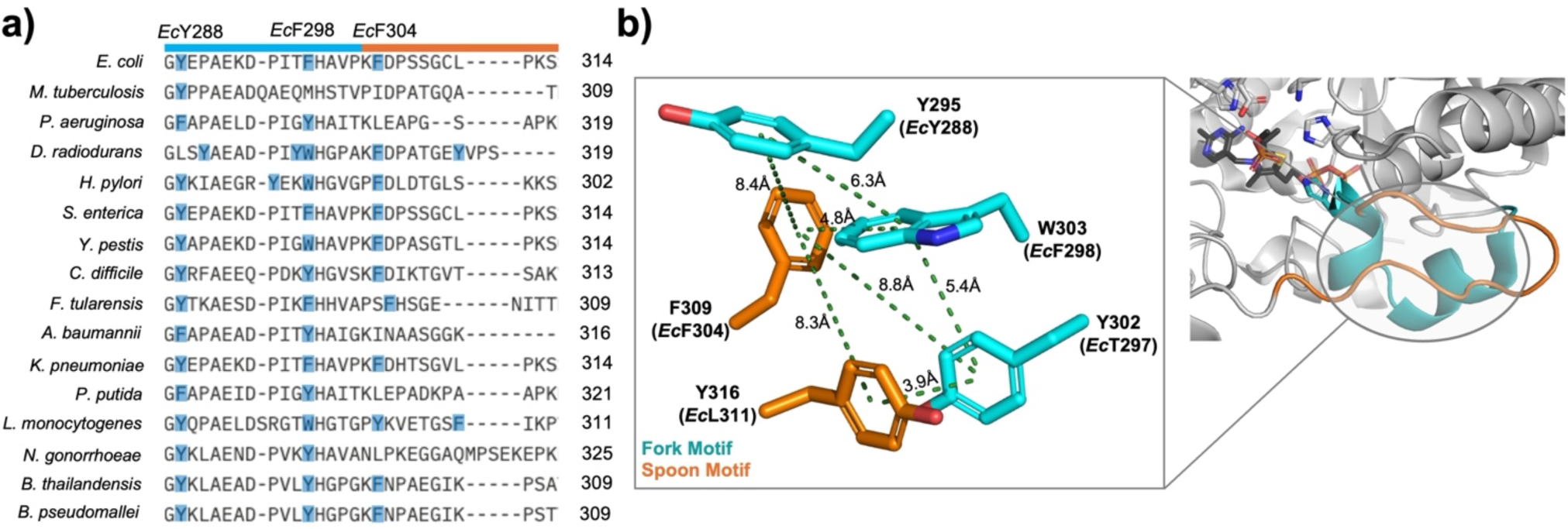
Sequence and structural alignments reveal conserved aromatic residues in the spoon and fork motifs. **a)** Sequence alignment of sixteen DXPS homologs reveals a conserved cluster of aromatic residues in the spoon (orange) and fork (cyan) mobile region, with the exception of *M. tuberculosis* (*Mtb*) DXPS which contains only 1 aromatic residue in this region). **b)** *Dr*DXPS aromatic residues in the spoon and fork motifs are in close proximity in the closed conformation.

Based on this analysis, we hypothesized that aromatic residues in the spoon and fork motifs (*Ec*Y288-F298-F304) assemble to form a cluster that stabilizes the closed conformation, promoting LThDP formation and subsequently preventing its activation (Figure S1c). Thus, we conducted computational and mutagenesis studies to test this hypothesis and elucidate how these aromatic residues influence DXPS function. For all variants, CD was used to detect changes in global secondary structure and stability. Kinetics of DXP formation, LThDP formation and persistence, and shifts in conformational equilibrium were investigated on DXPS variants to determine how the spoon and fork motifs participate in the ligand-gated DXPS mechanism.

### Aromatic cluster residues are predicted to be coupled to PLThDP and global DXPS motions in long-timescale simulations

As noted above, due to the link between conformation and catalysis, we hypothesized that the assembly of the aromatic cluster promotes the closed conformation and therefore would be correlated to reactivity of ThDP adducts within the active site. To predict how the dynamics of the aromatic cluster residues are involved in DXPS conformational transitions, we performed a long-timescale (23 µs) molecular dynamics simulation of PLThDP (derived from MAP) bound *Dr*DXPS with the Anton 2 supercomputer. Principal component analysis (PCA) was then performed on the resulting trajectory to characterize large-scale global dynamics of *Dr*DXPS and the PLThDP adduct derived from MAP (Figure 4a,b). The first principal component (PC1, computed by incorporating the *Dr*DXPS Cα and PLThDP heavy atoms) revealed regions of *Dr*DXPS known to be involved with open/close state transitions and LThDP stability undergo collective large-scale rearrangements. Specifically, the spoon and fork motifs (*Dr*DXPS 292−306 and 307−319), which contain the *Dr*DXPS aromatic cluster residues (*Dr*DXPS 295-302-303-309-316), are significantly displaced along PC1 (vertical green bars, Figure 4a), indicating their central role in *Dr*DXPS conformational transitions. The PC1 motion also shows that one of the bound PLThDP molecules undergoes substantial rearrangement, particularly in the MAP moiety of the PLThDP adduct within the protomer (PRO) B (Figure 4b, purple highlight). The active site network residues (*Dr*DXPS 51-101-304-423-430-434) involved in LThDP stabilization (vertical grey bars, Figure 4a) were all also found to contribute to the PC1 dynamic.^43^ Taken together, PC1 predicts large-scale *Dr*DXPS motions that are coupled to PLThDP motions. To investigate the extent to which aromatic cluster rearrangements are involved in the global PC1 *Dr*DXPS motion, we computed the correlation coefficient between individual aromatic pair distances and the first 10 principal components (Figure S2). Many of the individual aromatic pair distances show relatively strong correlations with the PC1 motion (Figure S2, orange/red blocks along the vertical above PC1), with correlations tending to deplete as the PC index increases (decreasing extent of global motion). Two of the strongest correlations with PC1 are found on opposing *Dr*DXPS monomers, involving PROA:309-316 (0.79) and PROB:295-316 (0.77). Two-dimensional free energy surfaces along the PC1 fluctuation and aromatic distances were constructed to appraise how these aromatic pairs interplay with *Dr*DXPS global transitions (Figure 4c). The aromatic pair PROA:309-316 was found to predominately inhabit a basin (∼4 Å pair distance and “negative” direction PC1 displacement, indicative of the closed state) that is confined by *Dr*DXPS conformational transitions along PC1 (Figure 4c, left). Although there was a brief transition to an extended distances, (>8 Å) the PROA:309-316 distance tended to remain smaller until the PC1 transition enabled access to larger distances, where the aromatic pair no longer displayed any strong conformational preference aside from a small meta-stable basin at a distance of ∼14 Å. On the opposing monomer, the aromatic pair PROB:295-316 showed less restrictive PC1 coupling features and larger distances (Figure 4c, right). The PROB:295-316 aromatic distance did not exhibit a strong conformational preference (broad basin of larger distances in the “negative” direction PC1 displacement) until the PC1 transition occurred. Motions in the “positive” PC1 direction constrained the PROB:295-316 pair to smaller distances, aside from a brief distance expansion near the midpoint of the PC1 motion. Although these two residues remained out of direct contact distance, their arrangement was still found to be coupled to *Dr*DXPS global motions. The free energy surface along the PROA:309-316 and PROB:295-316 distances indicates a “compensating” relationship between these aromatic pairs (Figure 4d). Indeed, the PROB:295-316 distance increased only when the PROA:309-316 residues were in close contact, and the PROA:309-316 distance only increased when the PROB:295-316 distance became relatively constrained. This type of relationship suggests that the aromatic clusters may be important to structural responses not only locally, but also across the DXPS dimer. Given the conservation of the aromatic residues in the spoon and fork motifs, as well as the predicted computational correlated motion between the *Dr*DXPS aromatic residues and the active site, we proceeded with a mutagenesis study to experimentally elucidate the role of the aromatic cluster in the DXPS mechanism.

**Figure 4:**
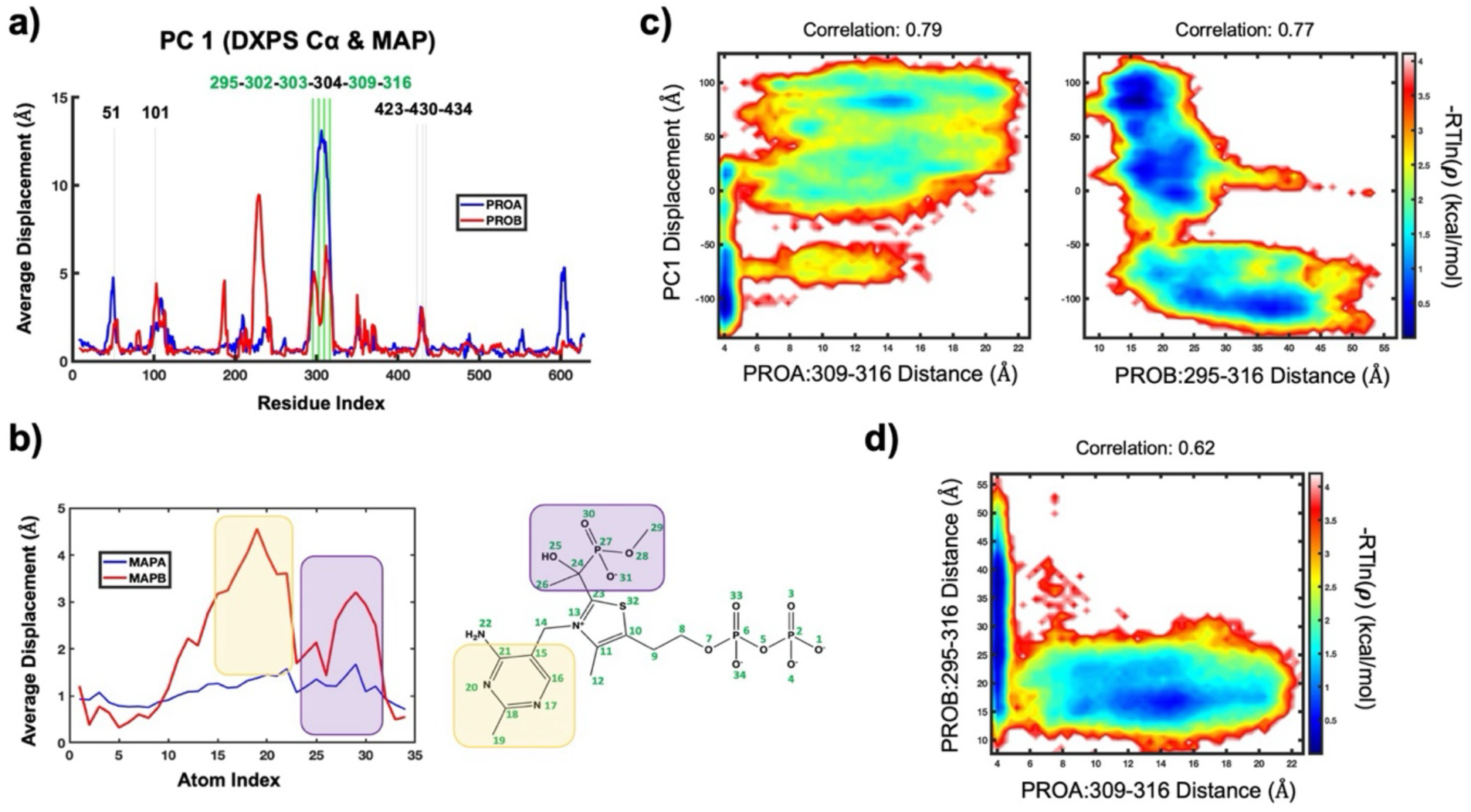
Aromatic clusters are predicted to be involved in PLThDP and global DXPS dynamics. The average displacement along the first principal component computed from the *Dr*DXPS Cα (shown in panel **a)** and the PLThDP (MAP) heavy atoms (shown in panel **b**). The vertical green bars in panel **a** are aligned to the aromatic cluster residues (*Dr*DXPS 295-302-303-309-316) and the previously reported active site network residues (*Dr*DXPS 51-101-304-423-430-434) are shown in gray bars. **c)** Free energy surfaces computed along the PC1 fluctuation and the PROA:309-316 distance (left) and PROB:295-316 distance (right). **d)** Free energy surface along the PROA:309-316 distance and PROB:295-316 distance.

### Substitution of Y288, F298, and F304 impairs DXP formation

Ala variants Y288A, F298A and F304A were generated to determine the effect of removing aromaticity and steric bulk. Additionally, conservative variants (Y288F, Y288Q, F298L, F304L) were generated to assess the impact of removing aromaticity or disrupting hydrogen bonding while maintaining steric bulk in the spoon and fork motifs. F298 and F304 were substituted with Leu based on the presence of leucine at these positions in other homologs (e.g., *P. aeruginosa* (*Pa*) DXPS, Figure 3a). Y288 was substituted with Phe or Gln to assess the importance of hydrogen bonding and aromaticity, respectively.

For each variant, the kinetic parameters for DXP formation were determined by varying pyruvate or D-GAP while maintaining the other substrate at a saturating concentration (Table 1 and Figure S3). We hypothesized that if Y288, F298 and F304 are necessary to maintain the closed conformation for LThDP formation, cluster disruption should impair catalysis. Indeed, Y288A, F298A and F304A variants all exhibited a ≥ 36-fold decrease in *k*_cat_ compared to WT DXPS, indicating an impaired rate-limiting step (Table 1 and Figures S3a,b,e,g). Interestingly, Y288A showed a 1173-fold decrease in *k*_cat_/*K*_m_^Pyruvate^ relative to WT DXPS, whereas *k*_cat_/*K*_m_^Pyr^ decreased 84-fold on F298A and 69-fold on F304A; this is consistent with a substantial increase in *K*_m_^Pyr^ on Y288A (*K*_m_^Pyruvate^ = 584 ± 40 μM) relative to F298A (*K*_m_^Pyruvate^ = 32 ± 1 μM) or F304A (*K*_m_^Pyruvate^ = 56 ± 2 μM). These results indicate that Y288 contributes to pyruvate binding despite its distance from the active site.

**Table 1:**
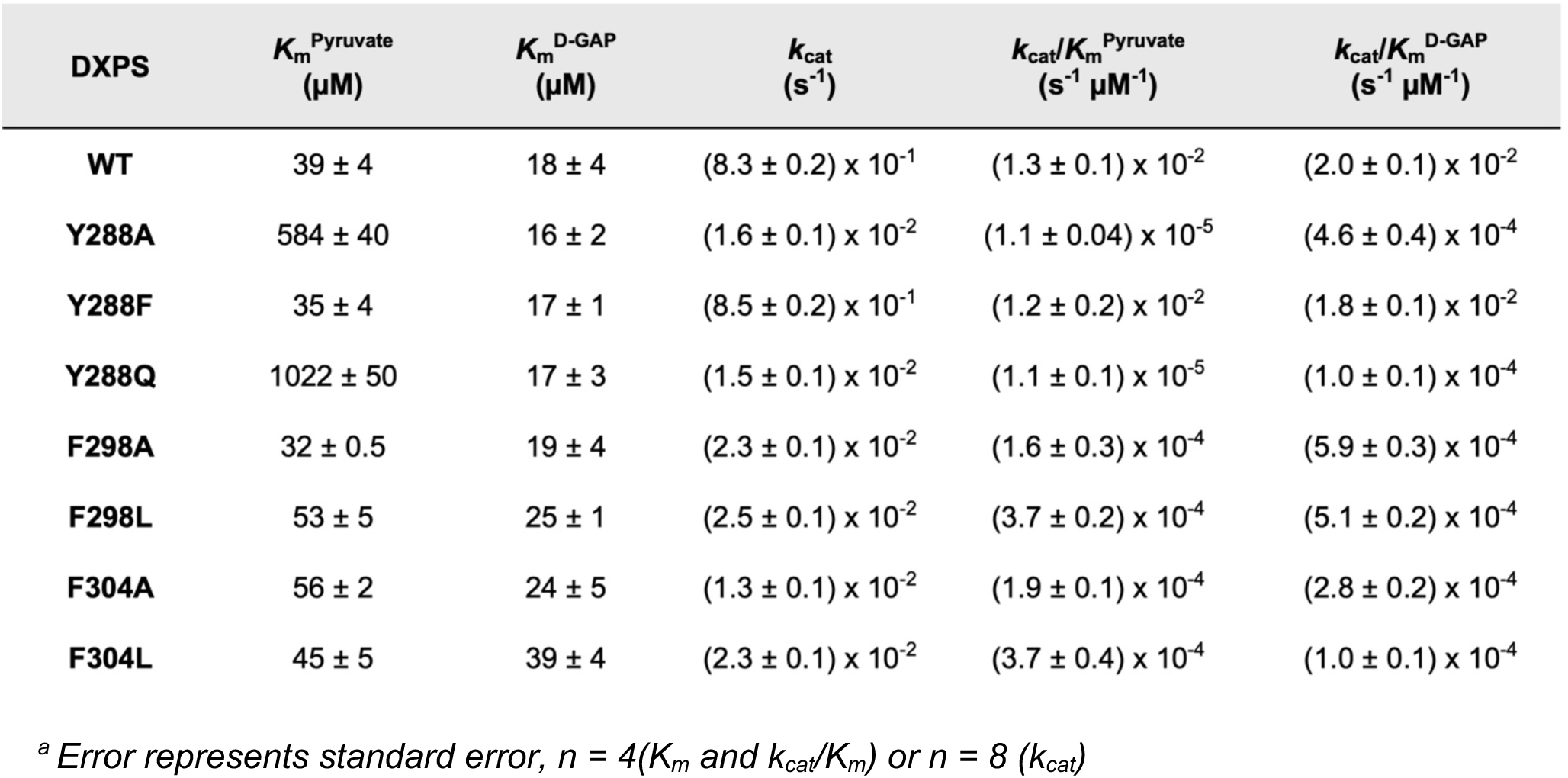
DXP formation kinetic characterization of WT and aromatic variant *Ec*DXPS.

To determine if these altered kinetic parameters were due to the loss of aromaticity or steric bulk in this region, we evaluated a series of conservative variants. We first asked whether loss of the hydroxyl or phenyl group of Y288 was contributing to the change in kinetic parameters observed for Y288A. The Y288F variant displayed kinetic parameters similar to WT DXPS, indicating that the hydroxyl group of Y288 is not involved in substrate binding or catalysis (Table 1 and Figures S3a,c). In contrast, the Y288Q variant displayed an 1249-fold decrease in *k*_cat_/*K*_m_^Pyruvate^ relative to WT DXPS, due to a 49-fold decrease in *k*_cat_ and a 26-fold increase in *K*_m_^Pyruvate^, indicating the importance of aromaticity at this position (Table 1 and Figures S3a,d). We next evaluated the importance of the phenyl groups of F298 and F304, by substituting these residues with Leu, based on the presence of leucine in other DXPS homologs at these positions. Both F298L and F304L displayed *K*_m_^Pyruvate^ similar to their alanine counterparts, each exhibiting a ≥ 33-fold reduction in *k*_cat_ (Table 1 and Figures S3a,f,h). In contrast to Y288Q, F298L and F304L maintained *K*_m_^Pyruvate^ comparable to the WT enzyme. With the exception of Y288F, which was comparable to WT DXPS, all other variants displayed *k*_cat_/*K*_m_^D-GAP^ 33- to 197-fold lower than WT enzyme, due to *k*_cat_ effects on F298L and F304L, and effects on both *K*_m_ and *k*_cat_ on Y288A/Q. Calculated *K*_m_^D-GAP^ for all variants were within ∼2-fold of *K*_m_^D-GAP^ for WT DXPS, suggesting that D-GAP binding is less influenced by assembly of the aromatic cluster in the spoon and fork motifs (Table 1 and Figure S3). Since Y288Q, F298L, and F304L displayed comparable kinetic parameters to their Ala counterparts and maintain steric bulk at these positions, we selected these three variants for subsequent experiments. CD experiments to assess global secondary structure and stability showed that Y288Q, F298L, and F304L are comparable to WT *Ec*DXPS (Figure S4 and Table S2). Thus, deviations in the behavior of variants relative to WT enzyme are unlikely to be a result of global enzymatic changes.

### Substitution of Y288, F298 or F304 alters the active site environment

CD is useful to assess perturbations in the DXPS active site environment, since it can detect changes to the CD profiles of 4’-AP and IP tautomers of the ThDP cofactor that are influenced by their local environment.^40,53–58^ ThDP bound to WT DXPS (E-ThDP) in the absence of substrate exists in the AP tautomer, which produces a CD signal with a λ_min_ at 320 nm (Figures 5a and S5a). The AP signal intensity is thought to arise from the charge transfer between the thiazolium and aminopyrimidine rings and is influenced by the active site environment; therefore, altering the active site environment manifests in deviations of the CD profile through signal shifts and/or changes in signal intensity.^15,43,53–59^ Given that substituting residues in the Y288-F298-F304 aromatic cluster impacted kinetic parameters, we hypothesized that these substitutions in the spoon and fork motifs may cause local conformational changes that would also alter the AP signal.

**Figure 5:**
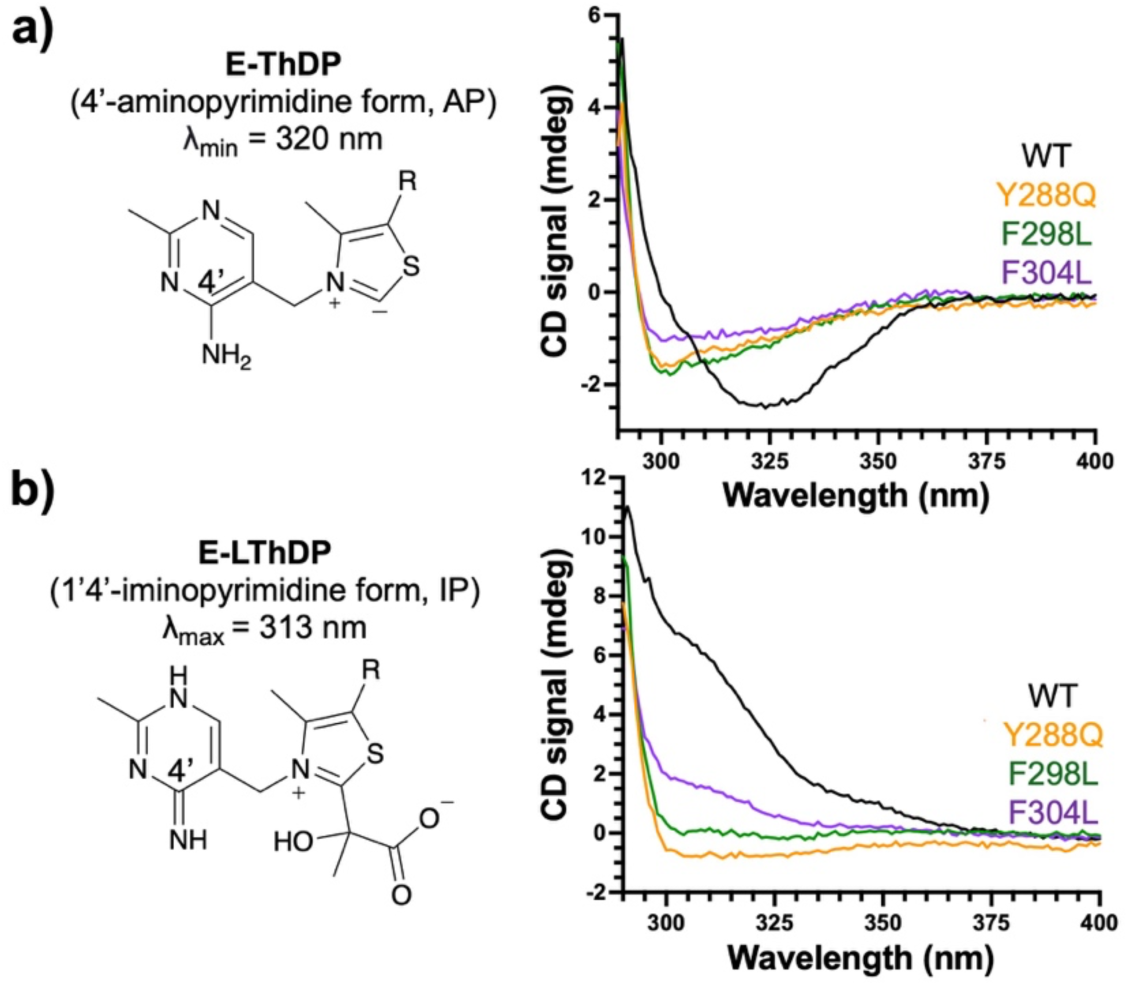
Y288Q, F298L and F304L display altered CD profiles. **a)** ThDP bound to WT DXPS exists as the AP tautomer having a signature CD profile with λ_min_ at 320 nm (AP structure on the left, CD_320_ signal in black on the right). All variants (30 µM) display shifts in the AP signal minima, shifted from 320 nm (WT DXPS) to 299-302 nm (right). **b)** LThDP forms upon reaction with pyruvate, producing the IP tautomer which inverts the CD signal to λ_max_ at 313 nm (IP structure on the left, CD_313_ signal in black on the right). All variants (30 µM) display lower IP signal maxima at 313 nm relative to WT DXPS when 5x*K*_m_^Pyruvate^ (200 µM for WT, 5 mM for Y288Q, 300 µM for F298L, and 200 µM for F304L) was added to form the LThDP intermediate (right). Replicate experiments can be found in Figure S5.

Compared to the WT enzyme, Y288Q, F298L, and F304L variants displayed shifts in the AP CD signal minima from 320 nm to 299-302 nm and lower negative CD signal intensities, indicating changes in the active site environment due to the removal of aromaticity in the mobile spoon and fork motifs (Figures 5a, S5a, and Table 2). Interestingly, these CD profiles are reminiscent of those reported previously for variants lacking H299 (e.g. H299A and H299N), the residue adjacent to F298 that is critical for catalysis and interacts with LThDP as part of the active site network on DXPS.^15,43^ Taken together, these results suggest disruption of the aromatic cluster within the spoon and fork motifs also alters the active site environment, possibly impeding the interaction of H299 with other active site network residues.

**Table 2:**
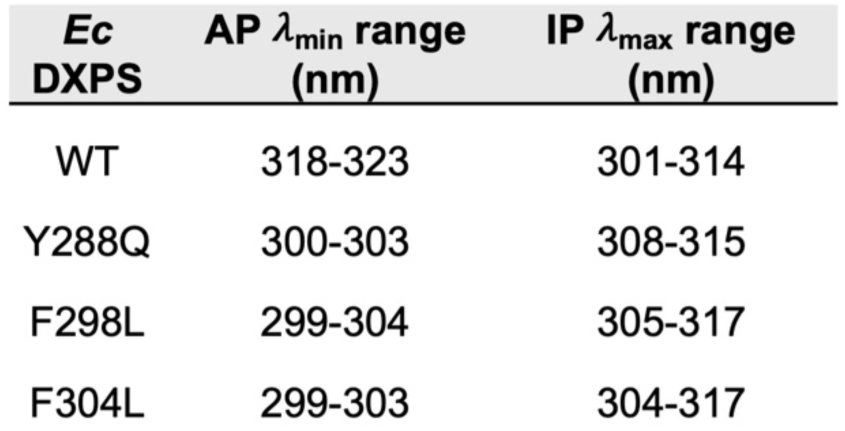
Active site architecture analysis by CD.

Upon formation of the LThDP intermediate on WT DXPS, the cofactor is converted to the IP tautomer, giving rise to an inverted CD signal with a λ_max_ at 313 nm (Figure 5b). As above, we predicted that changes to the active site environment caused by disruption of this aromatic cluster in the DXPS mobile regions would result in deviations from the WT IP signal. However, for each variant, we were unable to detect a positive CD signal in the presence of 100 µM pyruvate at 4 °C, conditions under which the CD_313_ signal corresponding to LThDP is detectable on WT DXPS. Thus, we increased the temperature to 25 °C and added pyruvate to 5x*K*_m_^Pyr^ for each variant. While a positive CD_313_ signal was observed on Y288Q, F298L, and F304L under these conditions, the signal intensities were significantly lower than that observed on WT DXPS (Figures 5b and S5b). The reduced signal intensities may indicate a variety of changes including slower LThDP accumulation, faster LThDP activation, and/or changes in LThDP molar ellipticity. Taken together, the observed changes in the AP and IP CD signatures for Y288Q, F298L and F304L suggest these aromatic residues within the spoon and fork motifs are necessary for proper organization of the active site cleft.

### Substitution of Y288, F298 or F304 impairs PLThDP formation

To assess how disruption of the aromatic cluster may impact LThDP formation, we used CD to observe phosphonolactylThDP (PLThDP) formation on Y288Q, F298L, and F304L upon MAP addition. MAP is a pyruvate mimic that forms a stable PLThDP adduct upon reaction with ThDP which is incapable of undergoing decarboxylation (Figure 6a). Similar to LThDP, the PLThDP adduct formed from MAP can be detected by CD (Figure S6a-d).^39^ Here, MAP was rapidly mixed with WT or variant DXPS using the stopped-flow apparatus, and the CD_313_ signal was monitored for 50 seconds at 10 °C. As expected, the PLThDP adduct was rapidly formed on WT DXPS (Figures 6b-d and S7e), as evidenced by the fast buildup of the CD signal at 313 nm. A CD_max_ of ∼ 4 mdeg was reached within 0.5 s and persisted for the duration of the 50 s experiment. In the absence of DXPS, reaction buffer mixed with 5 mM MAP showed no observable change in CD_313_ signal (Figures 6b-d and S7e). Interestingly, a negligible change in the CD_313_ signal was observed on Y288Q over 50 s (Figure 6b), while a slow increase in the CD_313_ signal was evident on F298L (Figure 6c) and F304L (Figure 6d) over this time period. Thus, PLThDP adduct formation was slow on these variants relative to WT DXPS, with the Y288Q variant displaying the most pronounced impairment in PLThDP formation over 50 s.

**Figure 6:**
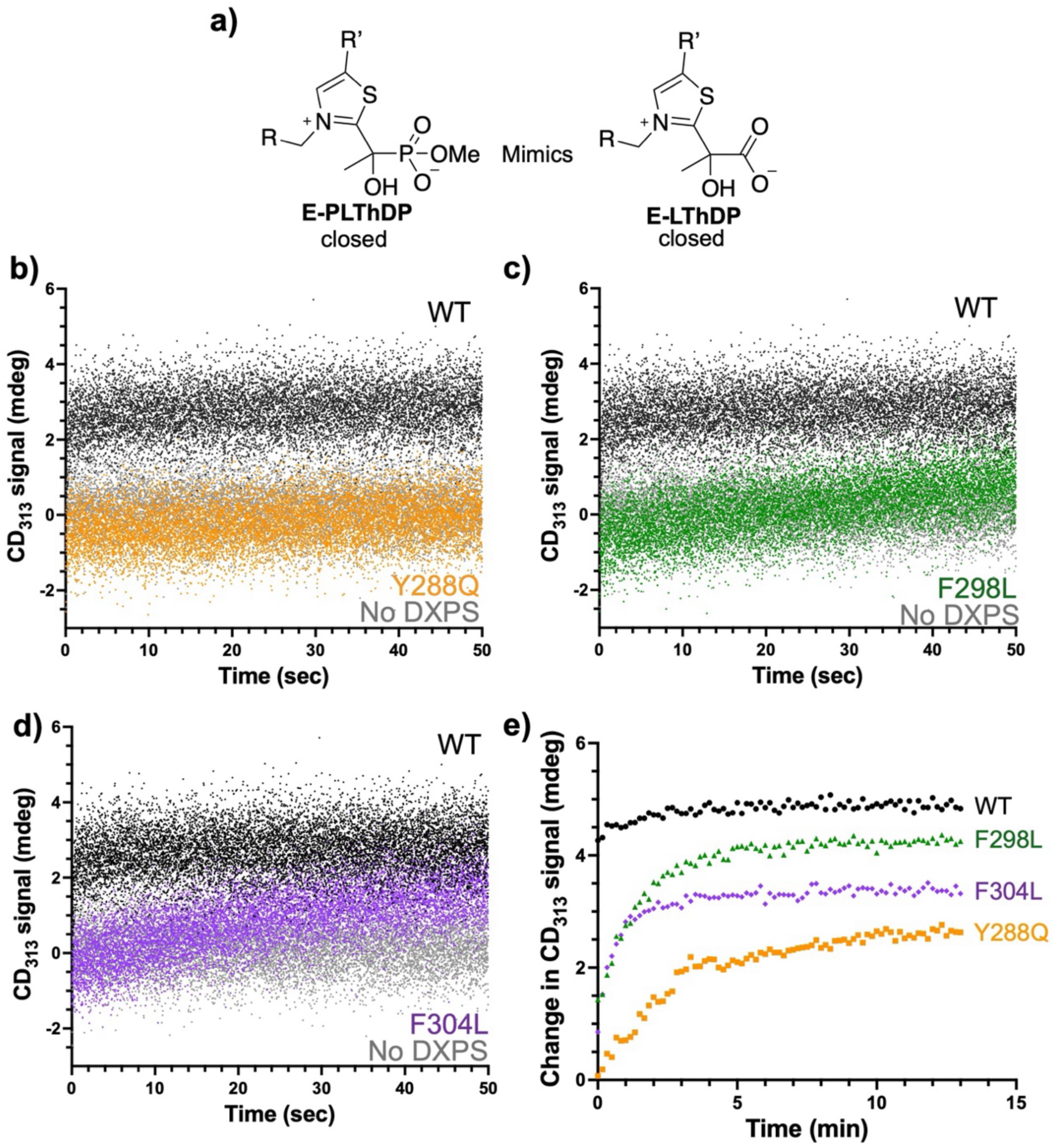
Disruption of aromatic cluster leads to slower PLThDP formation. **a)** DXPS binding to MAP produces PLThDP (left), a stable mimic of the natural LThDP intermediate (right). Time-resolved experiments monitoring of the PLThDP CD_313_ signal upon addition of MAP at 10 °C reveals limited PLThDP formation when the aromatic cluster is disrupted on Y288Q (**b**, orange), F298L (**c**, green), and F304L (**d**, purple) compared to WT DXPS (black) and a no DXPS (grey) control. **e**) Monitoring the CD_313_ signal at 10 °C over 13 min shows slower buildup of PLThDP when the aromatic cluster is disrupted, and reveals differences in CD_max_ for PLThDP on F298L, F304L and Y288Q under steady-state conditions. Signals are normalized to the AP signal in each case. Replicate experiments can be found in Figure S7.

A second CD experiment was performed to observe PLThDP formation over a longer time course and to determine CD_max_ on each variant. In this case, 5 mM MAP was added at 10 °C and the CD_313_ signal was measured every 10 s for 13 min. As expected, the PLThDP adduct formed quickly (within 1 minute) on WT DXPS under these conditions, reaching a CD_max_ of ∼ 3.6 mdeg (Figures 6e and S7a,b [normalized] and S7c [unnormalized]) Consistent with observations from time-resolved experiments, a more gradual rise in the CD_313_ signal was observed on Y288Q, F298L, and F304L. The maximal CD amplitude was observed by approximately 6 min on Y288Q, by 4 min on F298L, and by 2 min on F304L (Figures 6e and S7a,b[normalized]). Additionally, differences in the CD_max_ were observed for variants compared to WT DXPS (Figure S7c [unnormalized]), indicating differences in the PLThDP equilibrium concentration and/or the active site architecture. Taken together, these observations indicate PLThDP formation is hindered upon substitution of these aromatic residues, suggesting that disruption of the aromatic cluster impairs LThDP formation in the ligand-gated mechanism.

### Substitution of Y288, F298, or F304 shifts DXPS to an open conformation

LThDP formation and decarboxylation coincide with DXPS conformational changes to the closed and open forms, respectively (Figures 1 and 2).^13,33^ Based on our results showing slow accumulation of the PLThDP CD_313_ signal on Y288A, F298L and F304L variants (Figures 6 and S7), we reasoned that disruption of this aromatic cluster shifts the conformational equilibrium toward an open form which is thought to be less competent for PLThDP formation and stabilization. We previously demonstrated that limited trypsinolysis reports on the conformational equilibrium of *Ec*DXPS, offering insights into conformational shifts which are consistent with x-ray structural data.^13,15,33^ In the absence of substrate, WT DXPS exists in a conformational equilibrium between open and closed forms.^13,15,19^ Cleavage of *Ec*K227 occurs on either conformational form of full-length *Ec*DXPS, producing a 44 kDa peptide which is slowly converted to a 34 kDa peptide depending on DXPS conformation (Figures 7a,c and S8). Conversion of the 44 kDa peptide to the 34 kDa peptide on WT *Ec*DXPS occurs via cleavage at *Ec* K313, which is located in the spoon motif. Critically, K313 is protected from proteolysis in the closed conformation but susceptible to trypsinolysis in the open conformation. Addition of MAP leads to formation of a stable E-PLThDP complex and shifts the conformational equilibrium to the closed form in which K313 is protected from proteolysis. Thus, in the presence of MAP, the 44 kDa peptide is the major product and its conversion to the 34 kDa peptide occurs significantly more slowly relative to the (-) MAP control (Figures 7a,d).^15^ For DXPS variants that exist primarily in the open conformation, a more rapid accumulation of the 34 kDa peptide has been observed relative to WT DXPS. Thus, by measuring the cleavage pattern and rate of conversion of the 44 kDa peptide to the 34 kDa peptide, we can predict whether mutagenesis causes a shift in the conformational equilibrium.

**Figure 7:**
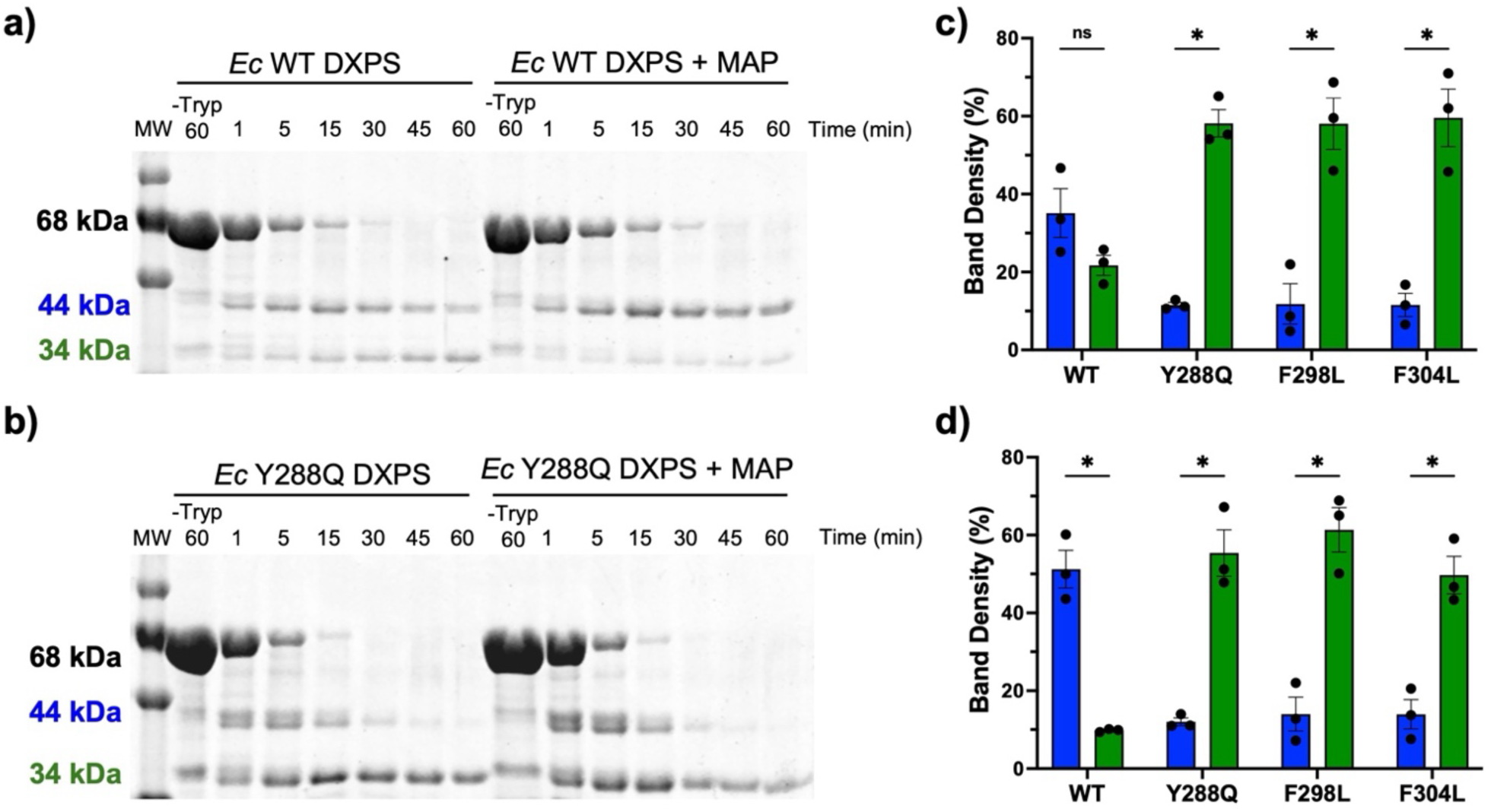
Substitution of Y288, F298 or F304 shifts the conformation to an open state. **a)** WT *Ec*DXPS in the absence (left) or presence (right) of 5 mM MAP. MAP stabilizes the closed conformation of WT *Ec*DXPS, in turn decreasing the rate of accumulation of the 34 kDa peptide. **b)** Trypsin digest time course samples from the representative Y288Q variant reveal accumulation of the 34 kDa band regardless of the addition of 5 mM MAP; **c)** Percent of total for 44 kDa (closed, blue) and 34 kDa (open, green) cleavage products 30 min after trypsin addition to WT or variant DXPS in absence MAP; **d)** Percent of total for 44 kDa (closed, blue) and 34 kDa (open, green) cleavage products 30 min after trypsin addition to WT or variant DXPS in presence MAP (5 mM); Error bars represent the standard error (n = 3), ns: not significant, *: p-value ≤ 0.05, **: p-value ≤ 0.01, ***: p-value ≤ 0.001.

Here, we used limited trypsinolysis to assess shifts in the conformational equilibrium of DXPS upon disruption of the aromatic cluster. Given our hypothesis that an intact aromatic cluster supports the closed conformation, we predicted that substitution of Y288, F298, or F304 would lead to more rapid accumulation of the 34 kDa peptide due to a shift to the open form. As expected, Y288Q, F298L, and F304L DXPS were more rapidly converted to the 34 kDa peptide (Figures 7b,c and S9), consistent with a larger portion of the enzyme population existing in an open conformation in each case. It is plausible that K313, the site of the second major cleavage event on WT *Ec*DXPS, becomes more accessible upon disruption of the aromatic cluster, resulting in a higher chance of cleavage at K313 prior to cleavage at K227 on full-length variants (Figure S8, top left). This alternate pathway is predicted to produce 33 and 34 kDa peptides which would contribute to more rapid conversion of full-length variants to smaller peptide fragments (33/34kDa) and less pronounced accumulation of the 44 kDa peptide (Figures 7b and S9,10). For all variants, additional peptide fragments near the 44 and 34 kDa peptides were evident, potentially arising from more pronounced cleavage at Lys and Arg residues nearby K227 and K313, which may become more accessible to trypsin in the open form of variants relative to WT *Ec*DXPS. Regardless, for all variants tested here, the peptide fragments near 44 kDa were nearly fully converted to the 34 kDa peptide by 60 minutes (Figures 7b, S9b-d, and S10), in contrast to WT DXPS (Figures 7a, S9a, and S10).

Since disruption of the aromatic cluster impairs PLThDP formation on each variant (Figures 6 and S7), we hypothesized that variants with disrupted aromatic clusters may not be protected from trypsinolysis upon incubation with MAP. Thus, we assessed the limited trypsinolysis profile of Y288Q, F298L, and F403L DXPS in the presence of MAP (5 mM). Each variant was incubated with MAP for 45 min to ensure ample time for PLThDP formation (Figures 6 and S7), followed by trypsinolysis over 60 min. Indeed, all three variants were rapidly digested to the 34 kDa peptide in the presence of MAP, similar to the (-) MAP control (Figures 7b,d, S9b-c, and S10). Thus, in contrast to WT DXPS, MAP does not have the effect to protect these variants from cleavage at K313, consistent with a shift to an open state in all three variants and an inability to stabilize PLThDP in a closed conformation.

As the global secondary structures and melting temperatures of these variants were comparable to WT DXPS (Figure S4 and Table S2), it is reasonable to suggest that the predicted changes in the conformational equilibrium are due to local structural changes within the spoon and fork motifs. These results, together with the observed slower formation of the PLThDP adduct on these variants, support the idea that disruption of the aromatic cluster shifts DXPS to an open conformation that is less competent for LThDP formation. Thus, disruption of the cluster may hinder conformational cycling required for efficient catalysis.

### F298L- and F304L-dependent LThDP activation

Based on the limited trypsinolysis results suggesting Y288Q, F298L, and F304L variants exist in a more open state thought to destabilize LThDP, we reasoned that disruption of the aromatic cluster should also have the effect to reduce LThDP persistence on DXPS.^43^ Stability of the CD_313_ LThDP (IP) signal can be assessed as a qualitative measure of LThDP persistence (Figure 8a).^19,43^ Thus, we investigated the stability of the CD_313_ signal on F298L and F304L DXPS; LThDP persistence on the Y288Q variant was not studied given the high *K*_m_^Pyr^ for Y288Q and lack of a pronounced positive CD_313_ signal corresponding to LThDP (Table 1, Figures 5b and S5b). Depletion of the CD_313_ LThDP signal on F298L and F304L DXPS was measured under anaerobic conditions where all solutions were degassed to avoid O_2_-mediated LThDP activation.^14^ Pyruvate was added to WT or variant DXPS to a final concentration of 5x*K*_m_^Pyr^, and a CD scan was immediately acquired (CD_313_ signal recorded within 2 min of adding pyruvate). As expected, a positive CD_313_ signal corresponding to the IP form of LThDP was observed. CD scans were then collected every 5 min at 25 °C until the signal depleted. This LThDP signal persisted on WT enzyme for ∼30 min before depleting (Table 2, Figures 8b and S11a). The IP signals for F298L and F304L DXPS reached a lower CD_max_ relative to the IP CD_max_ signal on WT DXPS, consistent with the lower signal amplitudes observed for these variants upon PLThDP formation (Table 2, Figures 6, 8c,d, S6, 7c, and S11b,c). In contrast to WT DXPS, the IP signals assigned to LThDP on F298L and F304L DXPS depleted rapidly with 42% and 32% reductions in the IP signals observed by 30 min for F298L and F304L variants, respectively (Figures 8e and S11d). The lower CD_max_ observed for the IP signal in the initial scan together with the more rapid decrease in IP signal intensity for F298L and F304L variants relative to WT DXPS is consistent with impaired LThDP formation and decreased LThDP persistence.

**Figure 8:**
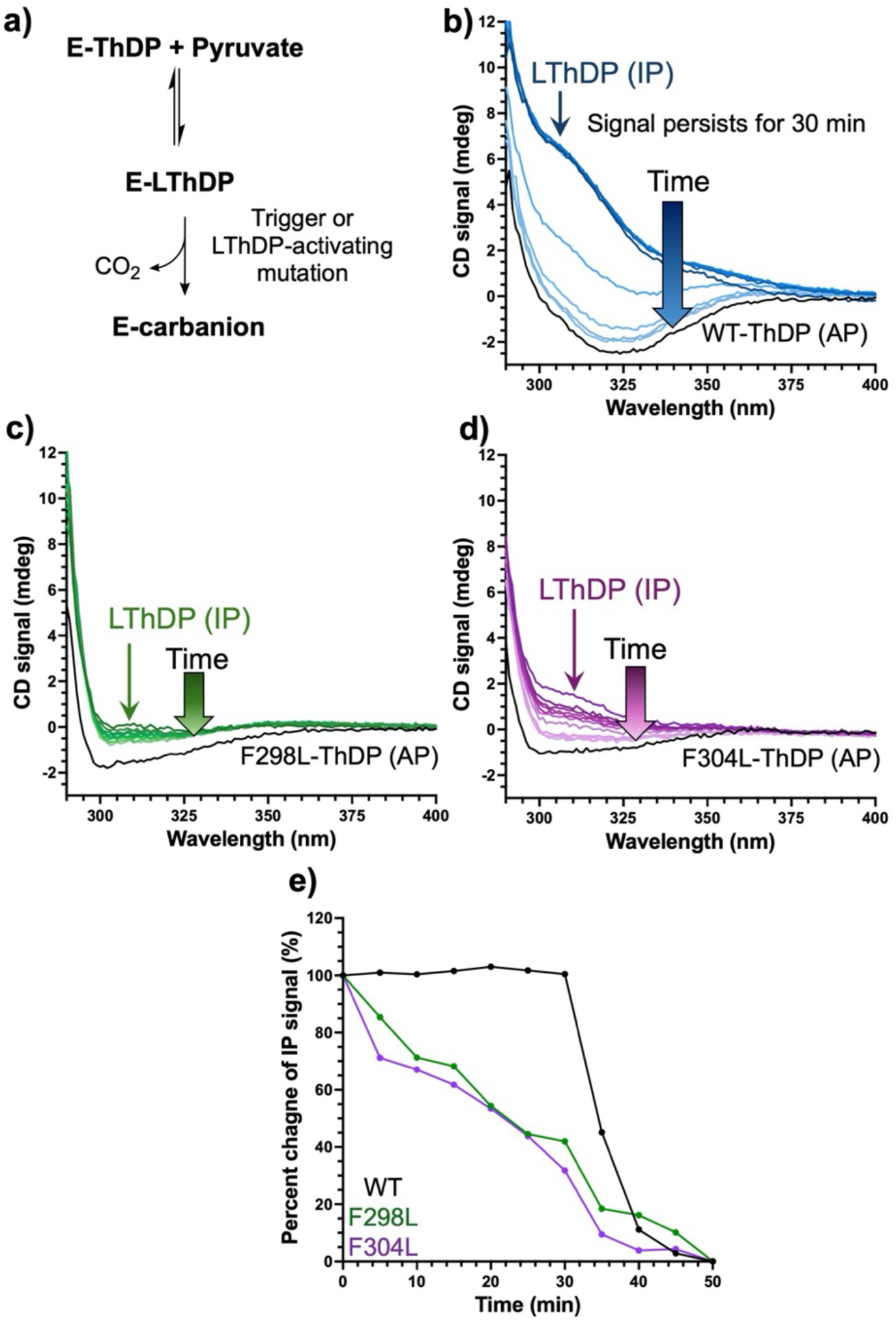
LThDP CD_313_ signal depletion upon disruption of the aromatic cluster. **a)** LThDP forms after pyruvate binding and persists in the absence of co-substrate or LThDP-activating mutations. **b)** Anaerobic monitoring of the CD_313_ LThDP signal on WT DXPS at 25 °C following addition of pyruvate (5x*K*_m_^Pyruvate^ = 200 μM) for 50 min. **c)** Anaerobic monitoring of the CD_313_ LThDP signal on F298L following addition of pyruvate (5x*K*_m_^Pyruvate^ = 300 μM) for 50 min. **d)** Anaerobic monitoring of the CD_313_ LThDP signal on F304L following addition of pyruvate (5x*K*_m_^Pyruvarte^ = 200 μM) for 50 min. e) Percent change of the total IP signal change in 60 min. Replicate data is shown in Figure S11.

### Detection of HEThDP upon the disruption of the aromatic cluster

We have previously shown that disruption of the DXPS active site network or addition of a trigger molecule leads to depletion of the CD_313_ IP signal which coincides with LThDP decarboxylation as confirmed by ^13^C NMR detection of HCO_3_^−^.^19,43^ Here, we used a similar approach to determine whether LThDP decarboxylation occurs more readily on Y288Q, F298L, and F304L variants. WT or variant DXPS (5 µM) was incubated with 500 µM [^13^C_3_]pyruvate anaerobically at ambient temperature (∼28 °C) for 20 min (Figure S13). For all variants, HCO_3_^−^ (163.03 ppm) was detected, as expected (Figures 9a, S14a). In addition, a new product peak at 24.77 ppm in the ^13^C NMR spectrum was observed in the presence of each variant, consistent with a two-carbon ^13^C-enriched species (Figures 9a and S14a). A product peak at 7.2 ppm in the ^1^H spectrum was also observed in the presence of each variant, consistent with the 6’-AP ring proton of a ThDP adduct (Figures 9b and S14b).^60,61^ Two-dimensional HSQC NMR of this product formed in the presence of F298L revealed correlations between ^1^H and ^13^C resonances at (5.42, 67.42 ppm) and (1.62, 24.77 ppm), with chemical shifts that are reminiscent of HEThDP.^51^ An authentic HEThDP standard was synthesized and compared to the unknown product. Indeed, upon analysis of the HEThDP standard by NMR under assay conditions, we observed 2D cross peaks at (5.42, 67.42 ppm) as well as cross peaks at (1.62, 24.77 ppm), assigned to C_2α_-CH and C_2α_-CH_3_ of the adduct, respectively (Figure S15). Additionally, a ^1^H resonance at 7.2 ppm was assigned to the HEThDP 6’-AP proton, providing further evidence for HEThDP as the unknown product. As expected, we observed an increase in the 7.2 ppm peak height upon addition of authentic HEThDP to the samples containing variant DXPS enzymes, confirming the presence of HEThDP in solution (Figure 9c). It is plausible that LThDP is readily decarboxylated in the active site, and then the enamine and/or HEThDP are released into solution. Alternatively, LThDP may be released from these variants, with decarboxylation occurring when the reaction is quenched by boiling.^62^ To learn if boiling promotes LThDP decarboxylation in solution, we modified the quench in an experiment using F298L DXPS. In lieu of quenching the reaction by boiling, we chilled samples after 20 min, and WT, and F298L DXPS was removed by centrifugation through a 10 kDa filter at 4 °C. Once again, the F298L sample showed evidence of HEThDP accumulation with ^13^C and ^1^H peaks at 24.77 and 7.2 ppm, respectively (Figure S16). This result suggests that minimal conversion of LThDP to HEThDP occurs during boiling, supporting a model in which aromatic cluster disruption promotes LThDP decarboxylation and ejection of the post-decarboxylation intermediate from the active site. Decreased affinity of ThDP adducts upon disruption of the aromatic cluster suggests these substitutions promote an open state that is distinct from the open state induced by disruption of the active site network, where a two carbon product was observed upon LThDP decarboxylation.^43^

**Figure 9:**
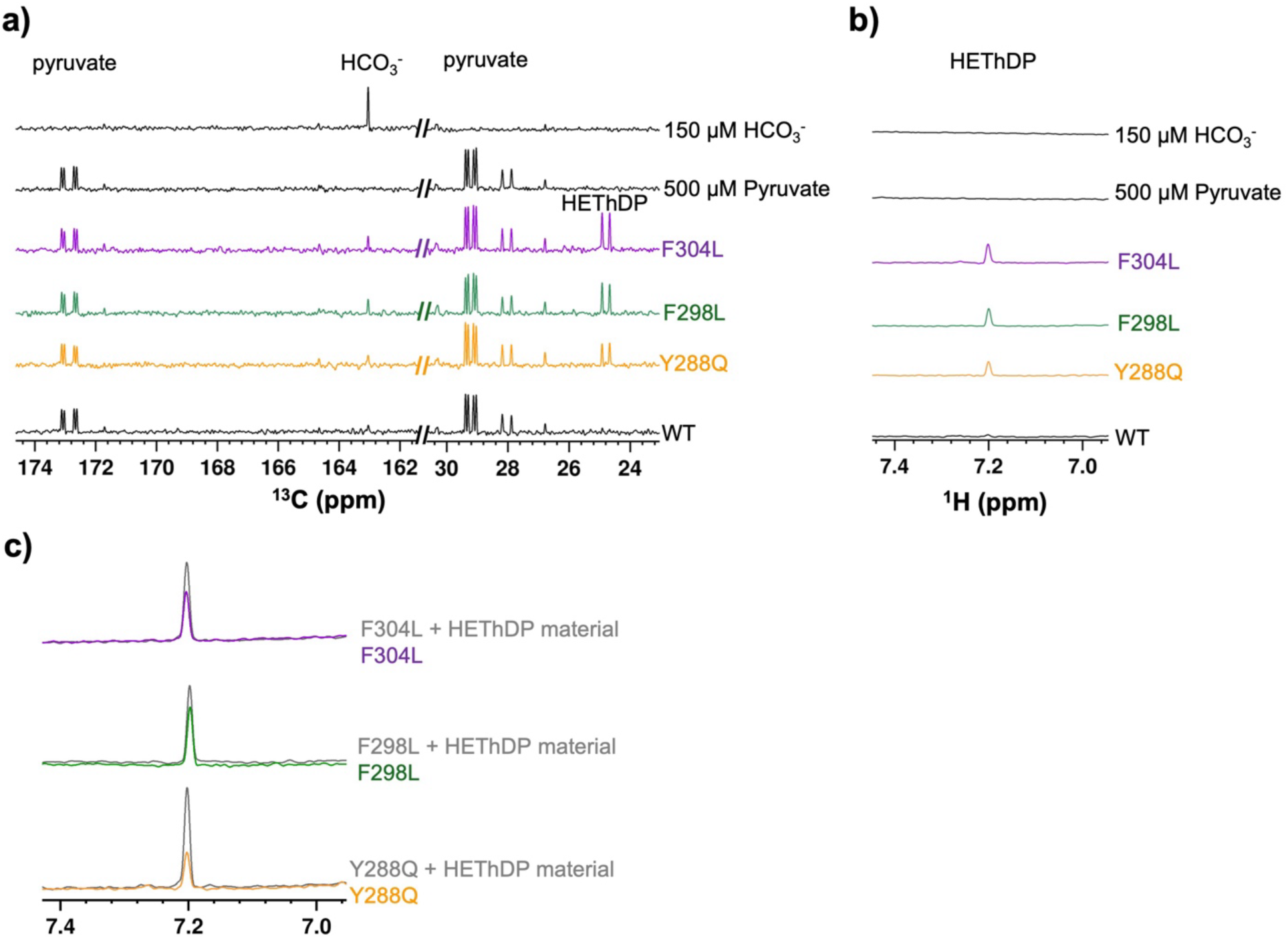
Evaluation of ThDP-intermediate loss for variant DXPS active sites. **a)** Bicarbonate was detected by ^13^C NMR upon disruption of the aromatic cluster by mutagenesis (purple, green, and orange), with minimal HCO_3_^−^ detected in the presence of WT DXPS (black). Spoon and fork variants Y288Q, F298L, and F304L generated a new product detected at 24.77 ppm consistent with HEThDP formation. **b)** ^1^H spectra acquired in the presence of Y288Q, F298L, and F304L variants exhibit a product peak at 7.2 ppm, consistent with HEThDP formation. Refer to Figure S14 for replicate NMR data. **c)** ^1^H NMR spectra showing the resonance at 7.2 ppm before (purple, green, and orange) and after (grey) addition of a HEThDP standard, confirming the accumulation of HEThDP in solution.

### H299A DXPS displays similar characteristics to aromatic variants

Given the reduced *k*_cat_ and similarity of CD signatures of Y288Q, F298L, and F304L DXPS variants relative to H299A DXPS, we hypothesized that the aromatic cluster functions to position this mobile catalytic histidine in the active site.^15^ To this end, we further examined the H299A variant to compare changes in intermediate formation and affinity using CD and NMR.^15,43^ Since H299A displays a reduced *k*_cat_, a shift to an open conformation, and an AP signal change relative to WT DXPS (shown here in Figures 10a and S17a as well as previously reported by our lab), we hypothesized H299A would also exhibit impaired PLThDP adduct formation.^15^ Indeed, H299A-catalyzed PLThDP formation reached equilibrium slowly, 5 min after addition of 5 mM MAP, in contrast to WT DXPS-catalyzed PLThDP formation which reached a maximum signal within 1 min (Figures 10b and S17b [normalized]). Similar to Y288Q, F298L, and F304L variants, the change in CD signal was lower on H299A compared to WT DXPS, with maximum intensities reaching ∼1.5 and ∼3.6 mdeg, respectively (Figures S7c and S17c [unnormalized]). This observed change in the H299A-PLThDP IP signal is consistent with either a lower equilibrium concentration of H299A-PLThDP relative to WT DXPS-PLThDP, and/or a change in the molar ellipticity of PLThDP on H299A (Figure S17c,d).

**Figure 10:**
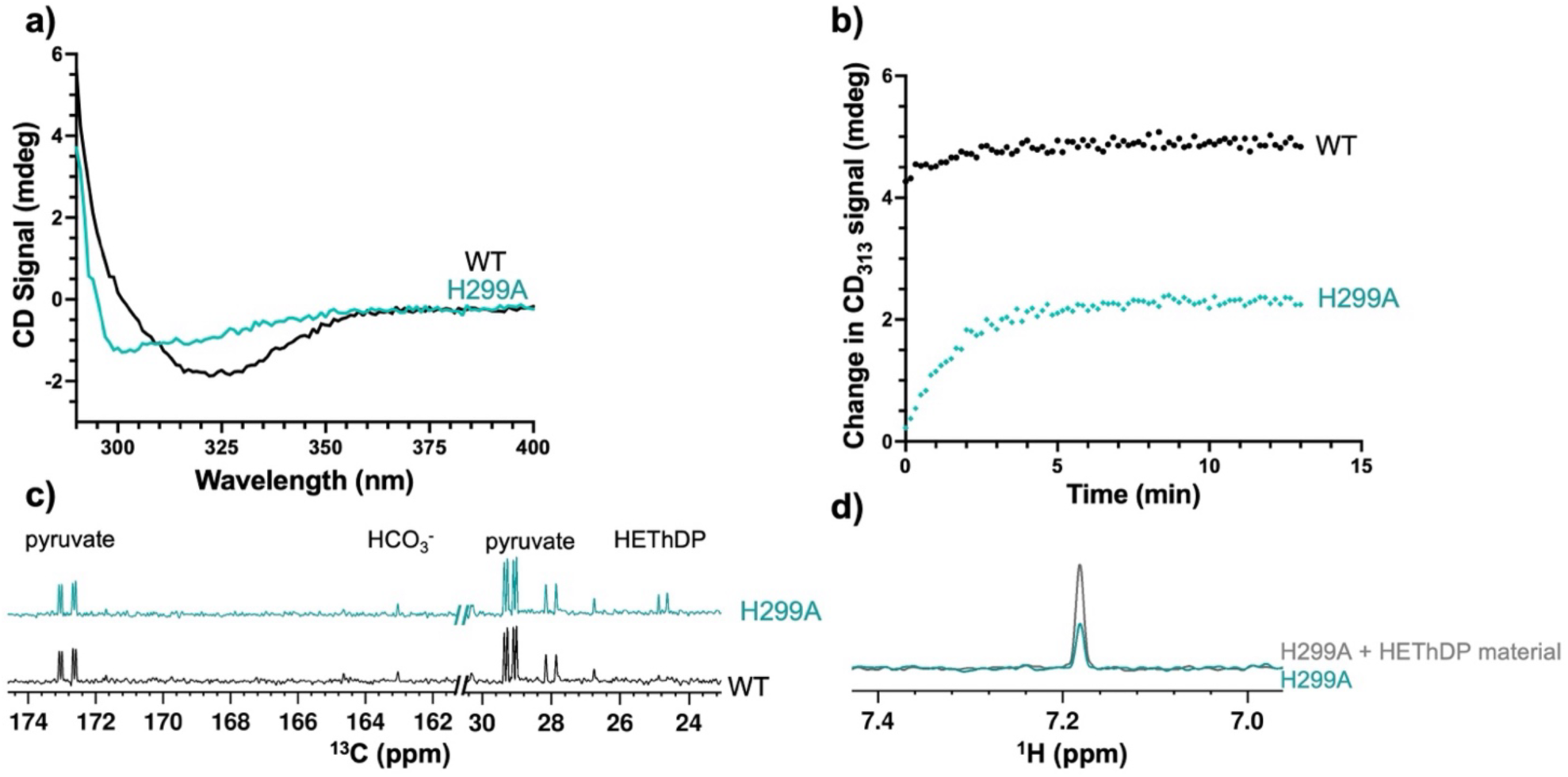
The H299A DXPS shares characteristics with Y288Q, F298L, and F304L DXPS. **a)** Representative CD spectra comparing WT DXPS (also shown in Figure 5e) and H299A AP signals under aerobic conditions; **b)** Representative CD time courses showing normalized PLThDP formation upon addition of 5 mM MAP under aerobic conditions, in the presence of WT or H299A DXPS. **c)** ^13^C NMR spectra acquired after incubation of 500 µM [^13^C_3_]pyruvate with WT or H299A DXPS. The peak at 24.77 ppm corresponding to the labeled hydroxyethyl group of HEThDP was detected in the H299A sample. **d)** The ^1^H spectrum of H299A DXPS (teal) shows a peak at ∼7.2 ppm correlating to the pyrimidine ring of HEThDP. Addition of a HEThDP standard (grey) results in an increased signal intensity at ∼7.2 ppm, supporting HEThDP accumulation in solution. All replicate data is shown on Figure S16.

We also hypothesized the H299A variant would exhibit lower affinity for ThDP adducts, similar to Y288Q, F298L, and F304L variants, detected as HEThDP accumulation in solution. Upon incubation of H299A DXPS with 500 µM [^13^C_3_]pyruvate as described above, we detected low concentrations of HCO_3_^−^ consistent with the slower intermediate formation and thus lower turnover (Figures 10c and S17e). Additionally, we detected ^13^C and ^1^H peaks at 24.77 and 7.2 ppm, respectively, indicating the presence of the HEThDP C2α-CH_3_ and 6’-AP protons, respectively (Figures 10c,d and S17e,f). Addition of the HEThDP standard resulted in an increase in the 7.2 ppm ^1^H peak corresponding to the 6’-AP proton of HEThDP, confirming accumulation of this species in solution. The presence of HEThDP in solution is consistent with H299A-mediated LThDP decarboxylation as well as decreased affinity for the enamine or HEThDP intermediates, similar to Y288Q, F298L, and F304L variants (Figure 10d). Overall, the comparable behavior of H299A and Y288Q, F298L, and F304L variants support the hypothesis that aromatic cluster assembly positions H299 within the active site upon binding of pyruvate.

## Discussion

The conformational dynamics of DXPS are tightly linked to catalysis. The positioning of the spoon and fork motifs changes dramatically along the DXPS reaction coordinate, and their movements are thought to dictate reactivity of intermediates in the active site.^13,15,33^ However, spoon and fork motifs are highly sequence-variable across DXPS homologs, and how these mobile regions support conformational and catalytic cycling on DXPS is incompletely understood.^33^ This work identified a conserved cluster of aromatic residues within the spoon and fork motifs that functions to stabilize the closed conformation, supporting an intact active site network for LThDP formation and persistence within the active site prior to ligand-induced LThDP decarboxylation on DXPS.

Spoon and fork motifs possessing three aromatic residues, noted in 9 of 16 sequences examined (including *Ec*DXPS), appear most common across homologs. It is possible that the variability in the number of aromatic residues may foretell mechanistic differences in the gated mechanism and/or substrate usage across DXPS homologs. *Mtb*DXPS contains only one aromatic residue in the spoon and fork motifs. Crystal structures of ThDP-bound *Dr*DXPS (5 aromatic residues) in its full-length or truncated forms show the spoon and fork motifs in the closed conformation, while the crystal structure of a truncated ThDP-bound *Mtb*DXPS shows a more open conformation as well as differently positioned active site network residues compared to *D*rDXPS.^9,26,33,35,43^ *Mtb*DXPS may be more conformationally flexible, which could, in theory, enable alternate product formation without the strict requirement for ligand gating.

MD simulation of *Dr*DXPS predicted correlations between aromatic cluster assembly, enzyme conformation and potentially active site reactivity, and suggested that the proximity of aromatic residues to one another is coupled to global structural changes. Interestingly, the distance of PROA309-316 (monomer A) was inversely related to PROB295-316 (monomer B) for the duration of the simulation; this is consistent with half-sites activity where only one monomer at a time maintains the closed conformation. Conformational control of active site reactivity is also observed in the E1 subunit of PDH.^57^

Each residue of the aromatic cluster is important for catalysis, with reductions in *k*_cat_ and slow PLThDP formation observed with all cluster variants, indicating the importance of cluster assembly in the rate-limiting step of LThDP formation on *Ec*DXPS. Interestingly, despite F298 and F304 being in close proximity to H299 which is critical for pyruvate binding and catalysis, they do not appear to contribute to pyruvate binding directly.^15^ In contrast, Y288 is critical for both substrate binding and catalysis. Of note, Y288 is also the most highly conserved aromatic residue in these mobile regions of DXPS homologs, suggesting it is important for DXP formation across species.

Given that conformational dynamics and catalysis are closely linked, it is not surprising that disruption of the cluster causes a shift to an open state which influences active site architecture and catalysis.^15,43^ Notably, the AP signals corresponding to ThDP bound to Y288Q, F298L, or F304L variants resemble the AP signal on H299A DXPS, which directly disrupts the active site network and causes a shift to the open state.^15,43^ It follows that disrupting the aromatic cluster should preclude assembly of an intact active site network by preventing optimal positioning of H299 to stabilize LThDP.

Indeed, all variants exist primarily in an open state, and disruption of the cluster destabilizes LThDP in each case, consistent with our findings that the closed state with an intact active site network is critical for stabilization of the E-LThDP complex.^43^ Our CD results showed that the IP CD_313_ signal did not fully deplete to re-establish the AP signal, likely due to signal degeneracy arising from newly established equilibria of various DXPS-bound ThDP adducts and/or CD active molecules.^42,53,55,57,61,64,65^

Interestingly, HEThDP was detected as the main product in addition to HCO_3_^−^ upon disruption of the aromatic cluster in the presence of pyruvate, indicating that disruption of the cluster also alters the active site in a distinct manner that results in intermediate ejection from the active site. As expected, HEThDP also accumulates in solution upon disruption of the active site network via substitution of H299 (H299A), consistent with the finding that H299A kinetics and CD profile are similar to Y288Q, F298L, and F304L; this supports the notion that disruption of the aromatic cluster effectively removes H299 from the active site, lowering the affinity for post-decarboxylation intermediates. It remains unclear which ThDP adduct is released from variant active sites – HEThDP or the enamine following DXPS-dependent LThDP decarboxylation. The inability to detect HEThDP in solution by CD after removal of enzyme may be due to low molar ellipticity of HEThDP in solution relative to its enzyme-bound form.^53,61^ Alternatively, if the enamine is released from the active site, its protonation in solution would result in racemic HEThDP, which is undetectable by CD.

This study reveals a new link between conformational dynamics and active site function. Previous studies evaluating *Ec*R99A, D427A, and H431A (active site network variants) revealed moderate differences in activity compared to WT DXPS, differences in the shift to open states, and generation of a 2-carbon product.^43^ In contrast, the present study demonstrates that disrupting the aromatic cluster not only dramatically reduces catalytic turnover and LThDP persistence, but also promotes release of a post-decarboxylation intermediate, likely due to improper positioning of H299. This difference signifies a distinct open state promoted by disruption of the aromatic cluster, that reduces affinity for ThDP adducts. While Y288, F298, and F304 do not directly interact with ThDP or intermediates, our results indicate this aromatic cluster plays an important role in maintaining the DXPS closed conformation to preserve its affinity for intermediates until binding of co-substrate. The combined DXPS sequence alignment and experimental results are consistent with a model for the functional aromatic cluster in which the highly conserved Y288 residue is necessary for H299 positioning within the active site for pyruvate binding. Assembly of the aromatic cluster supports transition to the closed conformation, maintaining H299 within the active site for both LThDP formation and persistence.

We have considered the possibility that the number of aromatic residues may be correlated with the inherent flexibility of the spoon and fork motifs in the closed conformation; homologs with fewer aromatic residues (e.g., *Mtb* or *Pa*DXPS) may be more flexible in this region compared to those with three or more aromatic residues (e.g. *Ec* or *Dr*DXPS). Interestingly, substitution of *Ec*F304 with Leu (observed on *Pa*DXPS) has a dramatic effect on *Ec*DXPS mechanism. However, *Pa*DXPS maintains similar DXP forming activity to WT *Ec*DXPS. In addition, bisubstrate analog inhibitors that promote a closed conformation and display nM potency on *Ec*DXPS are µM inhibitors of *Pa*DXPS.^31,44^ Together, these findings suggest other unknown factors may dictate the nature of the stable, closed E-LThDP conformation on other DXPS species.^31^

This study revealed a functional aromatic cluster outside the active site in the mobile spoon and fork motifs of *Ec*DXPS, which supports efficient LThDP formation and persistence through maintenance of the closed conformation. Thus, the aromatic cluster supports conformational cycling which drives catalysis on *Ec*DXPS. These results are significant as this is the first study to learn how these largely sequence-variable mobile regions support intermediate formation and reactivity despite being distal to the active site.

Trigger binding coincides with the transition from the closed state to open state for LThDP decarboxylation. D-GAP and alternate aldehydes have been shown to induce decarboxylation, which hints towards a sensing mechanism of DXPS to respond to cellular cues.^19^ It is tempting to speculate that the number of aromatic residues within these mobile regions reflects species-specific protective mechanisms which determine the sensitivity of DXPS homologs to alternative triggers and therefore alternate product formation. Finally, results from this study may inform potential new DXPS inhibitor design, such as inhibitors interfering with conformational cycling by targeting the aromatic cluster to either stabilize the open or closed conformations. Additionally, as suggested above, this work may foretell deviations in alternative substrate sensing and/or usage on different DXPS homologs, which may enable the development of species-specific inhibition strategies.

## Conclusions

This study examines a new structural feature that supports conformational cycling in the unique ligand-gated mechanism of *Ec*DXPS. The results indicate that aromatic residues in the spoon and fork motifs are necessary for positioning H299 within the active site and maintaining the closed conformation required for LThDP persistence on the enzyme. While we still do not know how binding of a trigger molecule induces conformational changes and LThDP decarboxylation, the characterization of the *Ec*DXPS aromatic cluster in this work provides additional evidence for conformational cycling that is tightly linked to DXPS catalysis and the ligand-gated mechanism.

## Supporting Information

General methods, DXPS global confermation (Figure S1), Variant DXPS primer sequence and annealing temperature for site directed mutagenesis (Table S1), Correlation between aromatic cluster pairs and *Dr*DXPS simulation principal components (Figure S2), Aerobic DXP formation Michaelis Menten Curves (Figure S3), Secondary Structure and apparent melting temperature comparison of WT and variant DXPS (Figure S4), WT and variant DXPS apparent melting temperature (Table S2), Duplicate curves comparing WT and variant DXPS AP and IP ThDP tautomers using steady state CD signatures (Figure S5), WT and variant duplicate AP and PLThDP IP signatures (Figure S6), PLThDP formation on WT and variant DXPS (Figure S7), Trypsinolysis scheme (Figure S8), Accumulation of trypsin digest products (Figure S9), Raw triplicate trypsin digest gels (Figure S10), Clarified solution analysis upon WT and variant DXPS LThDP IP signal depletion methods, Duplicate LThDP persistence and clarified solution analysis of WT and variant DXPS using steady state CD (Figure S11), WT DXPS and dialysis buffer comparison of AP and LThDP IP tautomers by steady state CD (Figure S12), Observing time-resolved PLThDP formation at 25°C methods, Aromatic cluster disrupting variants show slower PLThDP at 25 °C (Figure S13), Observation of intermediate release from aromatic disrupting variants (Figure S14), NMR comparison of HEThDP standard to the F298L unknown product methods, Comparison of HEThDP and F298L unknown product by NMR (Figure S15), Discernment between pre- and post-decarboxylation intermediate by NMR (Figure S16), Duplicate data comparing H299A to aromatic removing variants (Figure S17), References

## Accession Codes

*D. radiodurans* DXPS Uniprot: Q9RUB5, NCBI: Q9RUB5; *E. coli* DXPS Uniprot: P77488, NCBI: Q0TKM1; *E. coli* MEP synthase Uniprot: P45568, NCBI: ATZ31749

## Author contributions

L.J.K and C.L.F.M. designed the study. L.J.K. performed all biochemical and biophysical experiments. A.M. designed, optimized, and assisted with 1D NMR acquisition, as well as performed 2D NMR analysis. N.D.S synthesized the HEThDP standard material. S.L.A. and H.L.W designed and performed all computational experiments. L.J.K. and C.L.F.M prepared the manuscript which all authors reviewed and gave approval of the final version.

## Acknowledgements

This manuscript is the result of funding in whole or in part by the National Institutes of Health (NIH). It is subject to the NIH Public Access Policy. Through acceptance of this federal funding, NIH has been given a right to make this manuscript publicly available in PubMed Central upon the Official Date of Publication, as defined by NIH. L.J.K, N.D.S., and C.L.F.M would like to thank the NIH’s support of the NMR facility which supported data acquired using the JHU-Pharmacology JEOL JNM-ECZL500R spectrometer through award number S10OD034217. The work performed by L.J.K, N.D.S, and C.L.F.M. was supported by NIH grants R01GM143810, T32-GM007445, and T32GM149382 at JHU. L.J.K and C.L.F.M would like to thank Eucolona M. Toci and Alicia A. DeColli for providing *Ec*H299A. S.L.A and H.L.W acknowledge Research Computing at the University of South Florida for providing services towards the completion of this study. S.L.A and H.L.W. would additionally like to thank the NIH’s support via R01GM116961 for the Pittsburgh Supercomputing Center (PSC) which allowed Anton 2 computer time (supported by grant MCB210018P) for this work. D.E. Shaw Research generously made the Anton 2 machine available at PSC. The content is solely the responsibility of the authors and does not necessarily represent the official views of the National Institutes of Health.

## Abbreviations

DXPS: 1-deoxy-D-xylulose 5-phosphate synthase
D-GAP: D-glyceradehyde-3-phosphate
MAP: methyl acetylphophonate
ThDP: thiamin diphosphate
LThDP: C2⍺-lactyl-thiamin diphosphate
HEThDP: 2-(1-hydroxyethyl)thiamine diphosphate
PLThDP: C2⍺-phosphonolactylthiamin diphosphate
HDX-MS: hydrogen-dueterium mass spectrometry
AP: aminopyrimidine tautomer of ThDP
IP: 1’,4’-iminopyrimidine tautomer of ThDP

## Supporting Information

**Figure S1:**
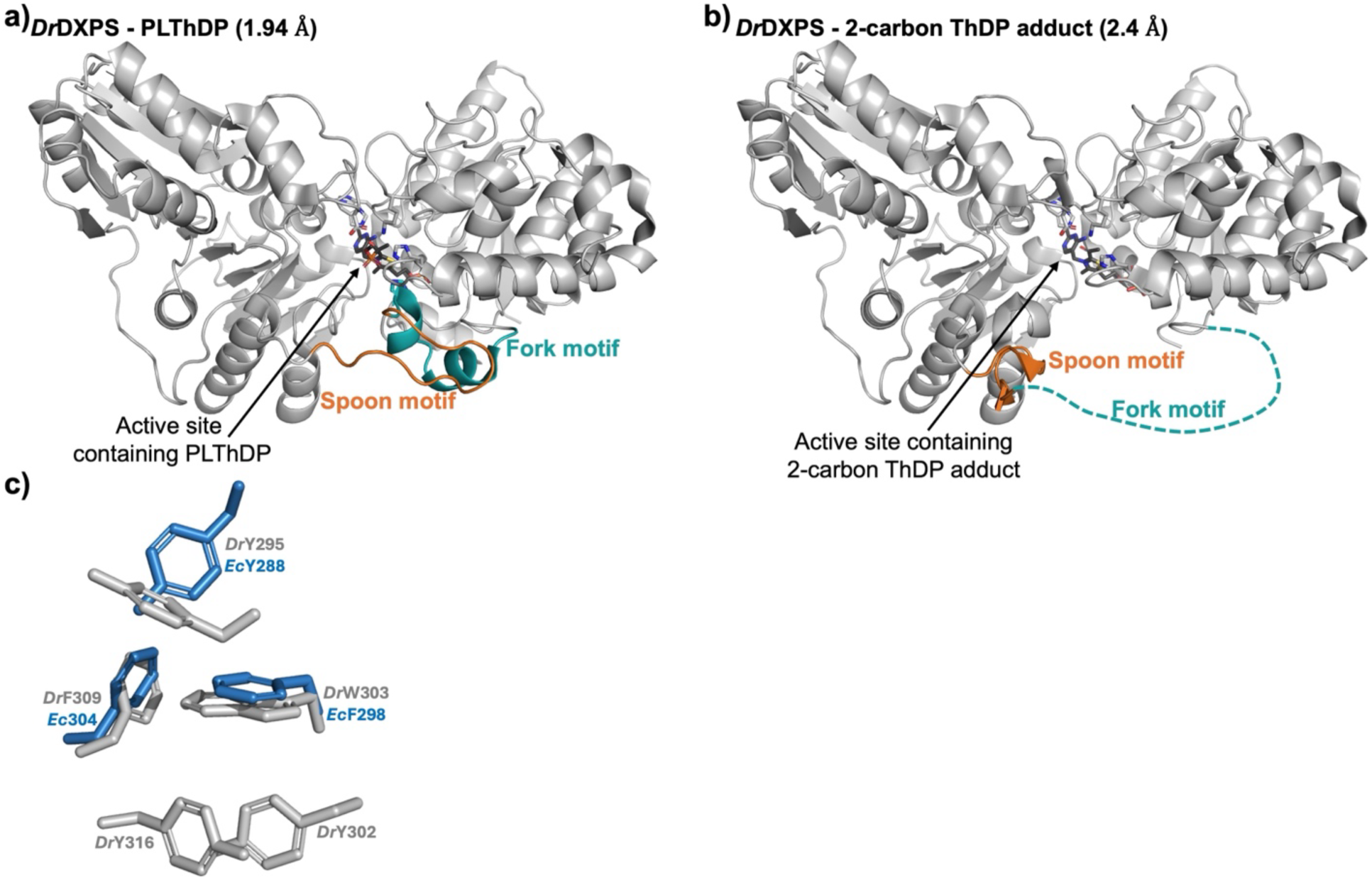
DXPS undergoes global conformational changes during catalysis, two of which have been determined previously: the closed (a, PBDID: 6ouv) and open (b, PBDID: 6ouw).^1^ c) A structure alignment of the *Dr* (grey) crystal structure and *Ec* (blue) AlphaFold model reveals spatial alignment of the spoon and fork aromatic residue clusters.

**Table S1:**
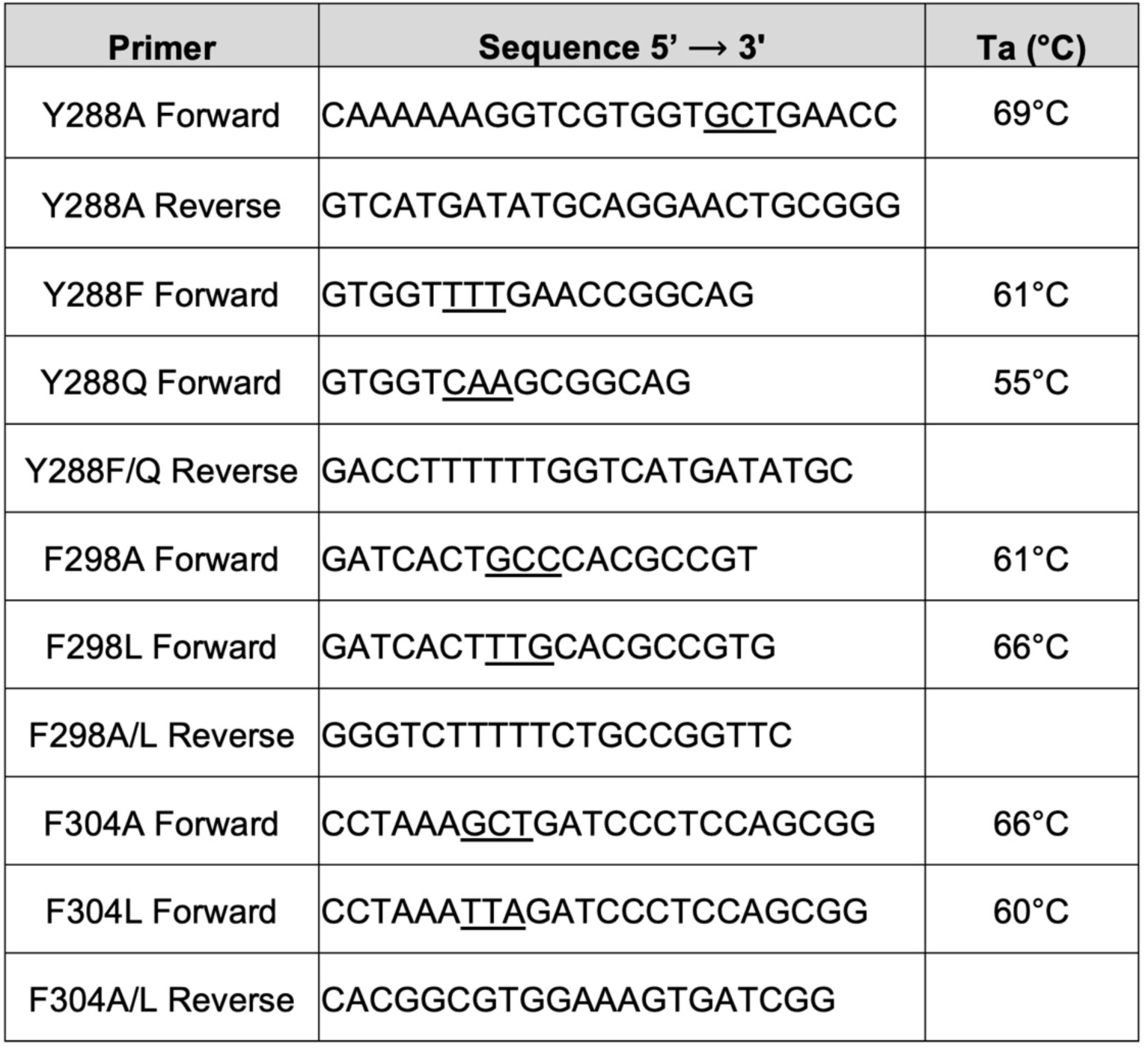
Variant DXPS primer sequence and annealing temperature for site directed mutagenesis.

**Figure S2:**
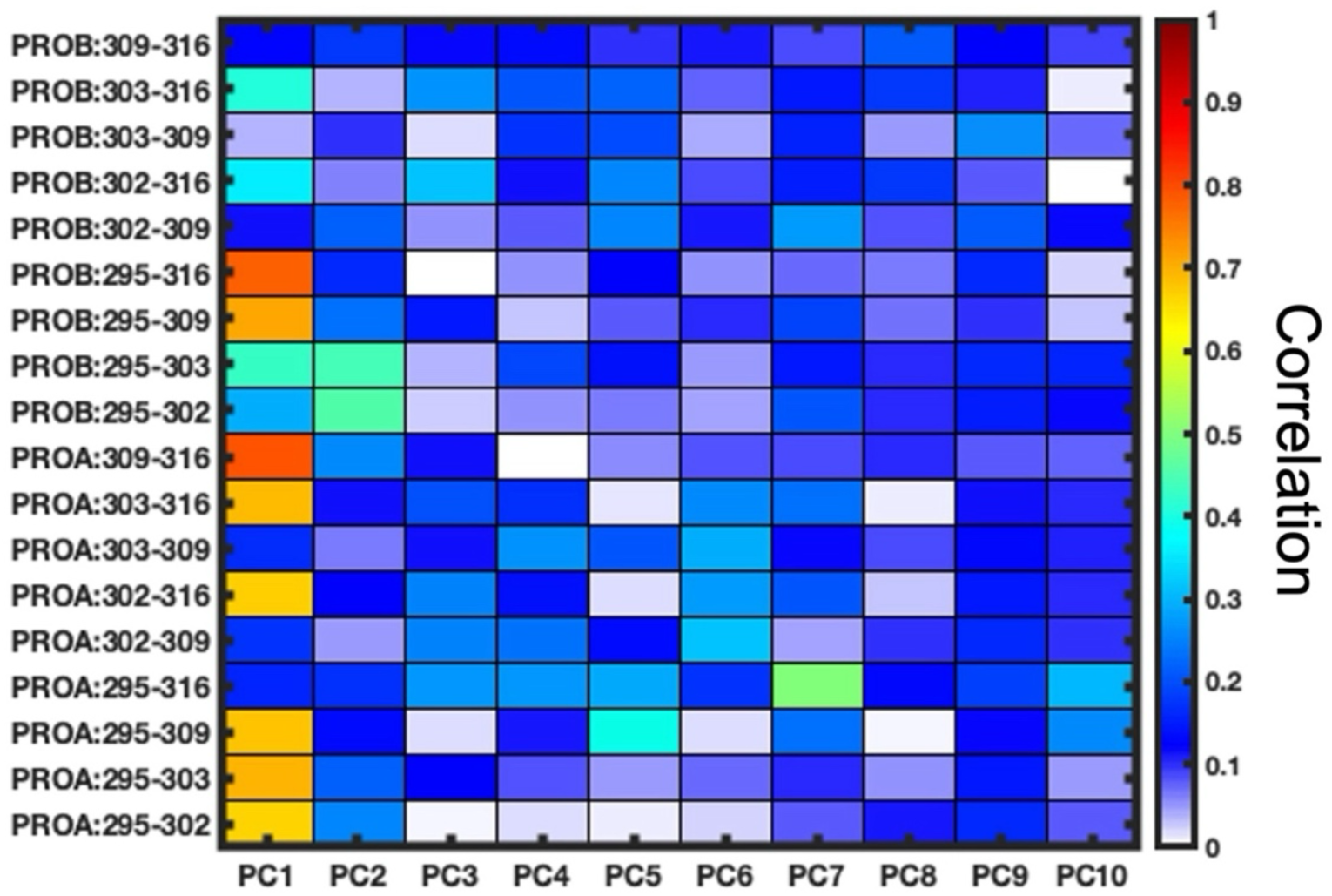
Aromatic cluster dynamics are correlated to global DXPS dynamics. The correlation values were computed between the aromatic cluster pair distances and the principal components.

**Figure S3:**
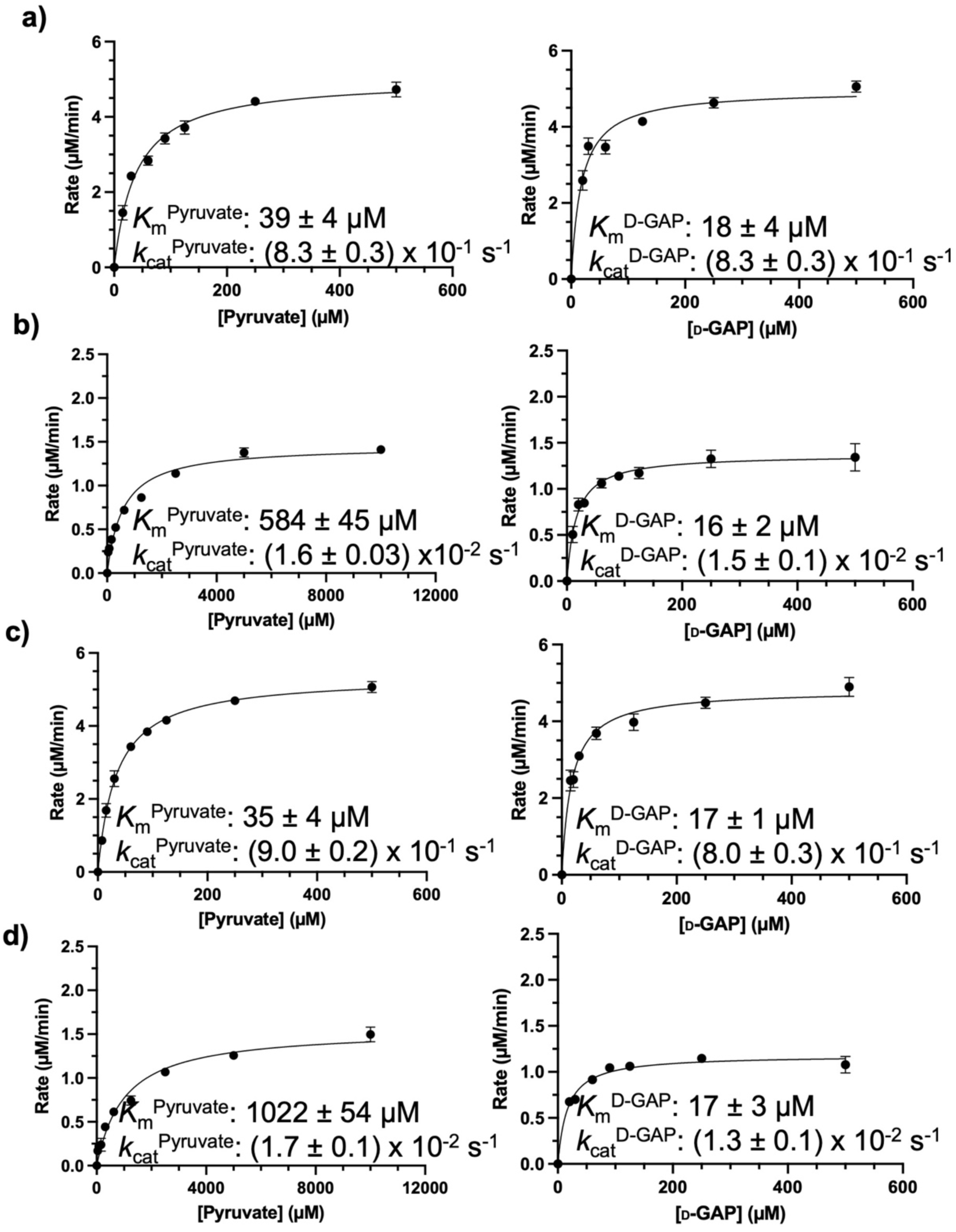
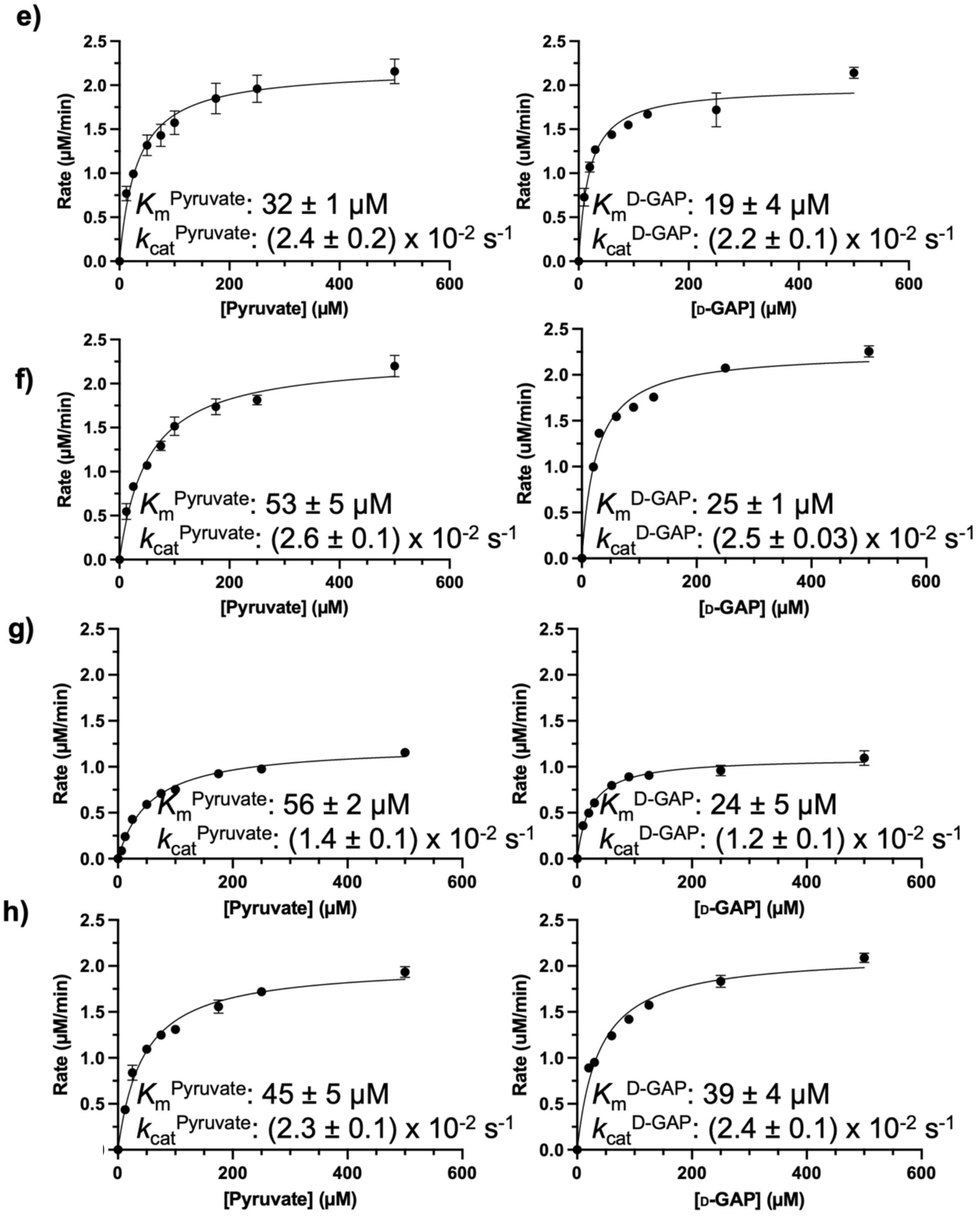
Determination of kinetic parameters for DXP formation on WT and variant DXPS. Substrate concentrations are varied (pyruvate, left; D-GAP, right) under aerobic conditions on WT (a), Y288A (b), Y288F (c), Y288Q (d), F298A (e), F298L (f), F304A (g), and F304L (h). Error bars represent the standard error of the mean. Errors in (*K*_m_) and (*k*_cat_) represent standard error determined from 4 experiments.

**Figure S4:**
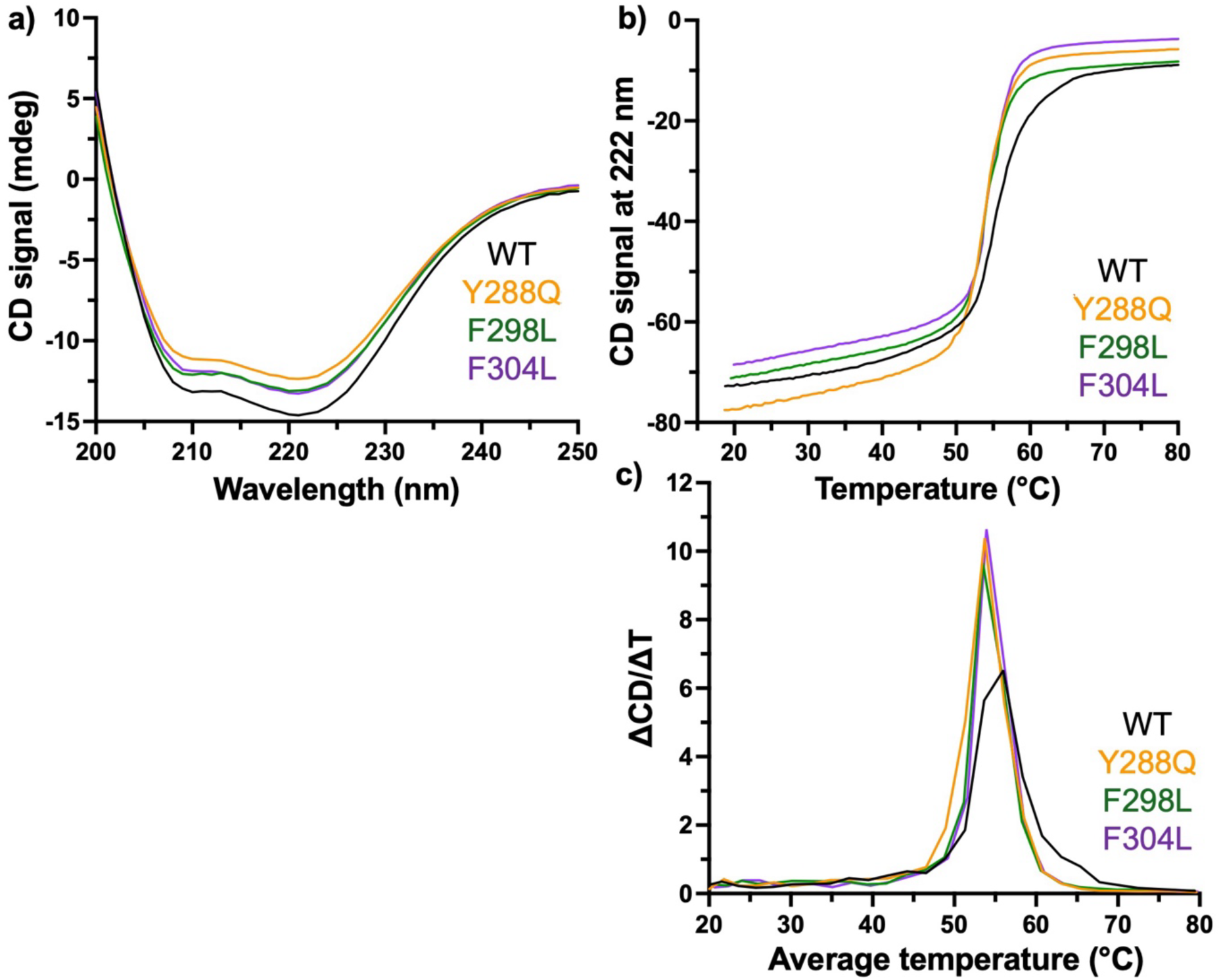
Representative secondary structure (a) and thermal stability (b,c) curves of WT and variant DXPS measured by CD (n = 3). a) The ⍺ helical nature is maintained on variant DXPS. b) CD signal was measured from 20–80 °C to determine the thermal stability. c) The first derivative was calculated to estimate the apparent melting temperature (thermal denaturation was not reversible) using the maximum of each curve.

**Table S2:** WT and variant DXPS apparent melting temperature.

| *Ec* DXPS | $appT_m$ (°C) |
| --- | --- |
| WT | 55.0 ± 0.5 |
| Y288Q | 54.0 ± 0.5 |
| F298L | 53.2 ± 0.4 |
| F304L | 54.3 ± 0.2 |

Error represents standard error, n = 3

**Figure S5:**
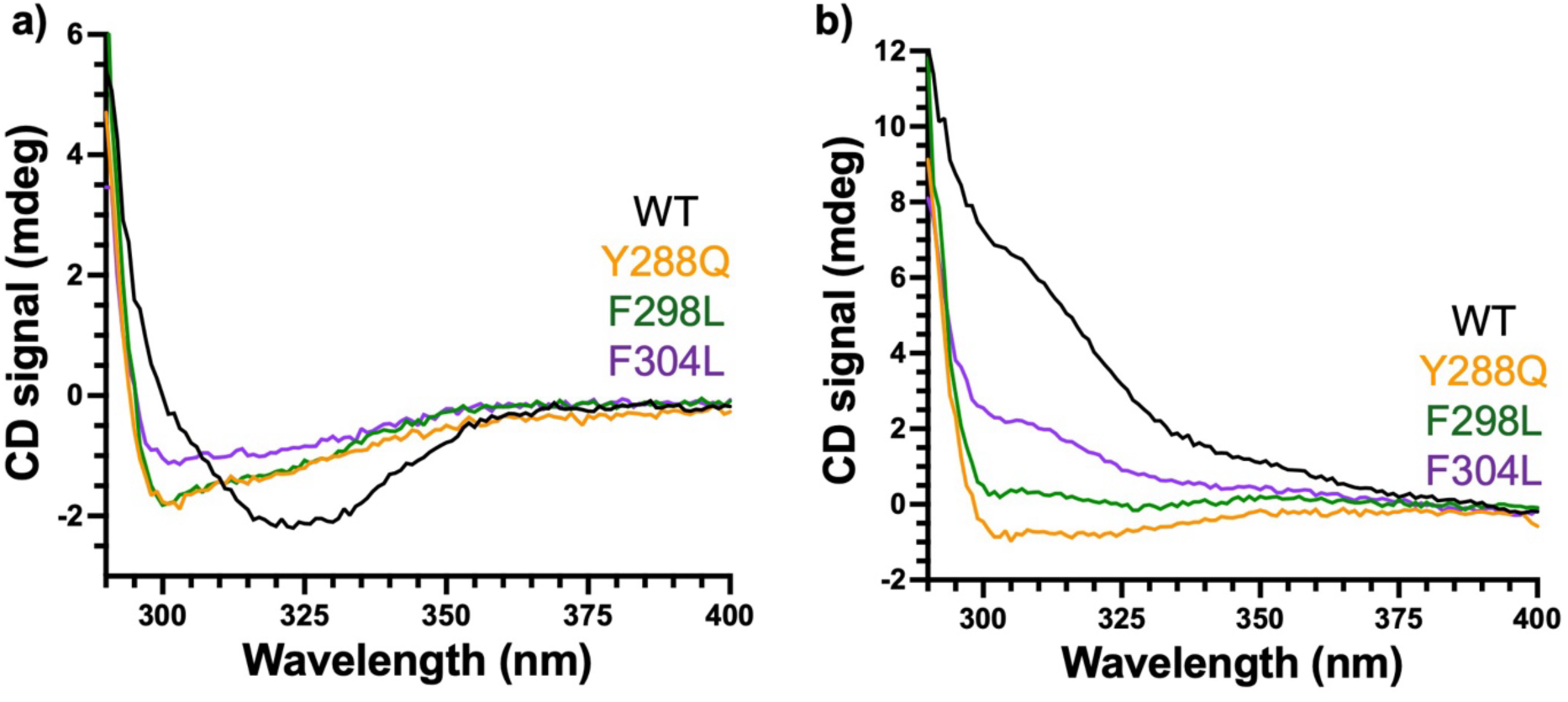
Replicate CD curves of WT and variant DXPS AP (a) and LThDP IP (b) signals. LThDP was formed upon the addition of pyruvate (final concentration of 5x*K*_m_^Pyr^) in each case.

**Figure S6:**
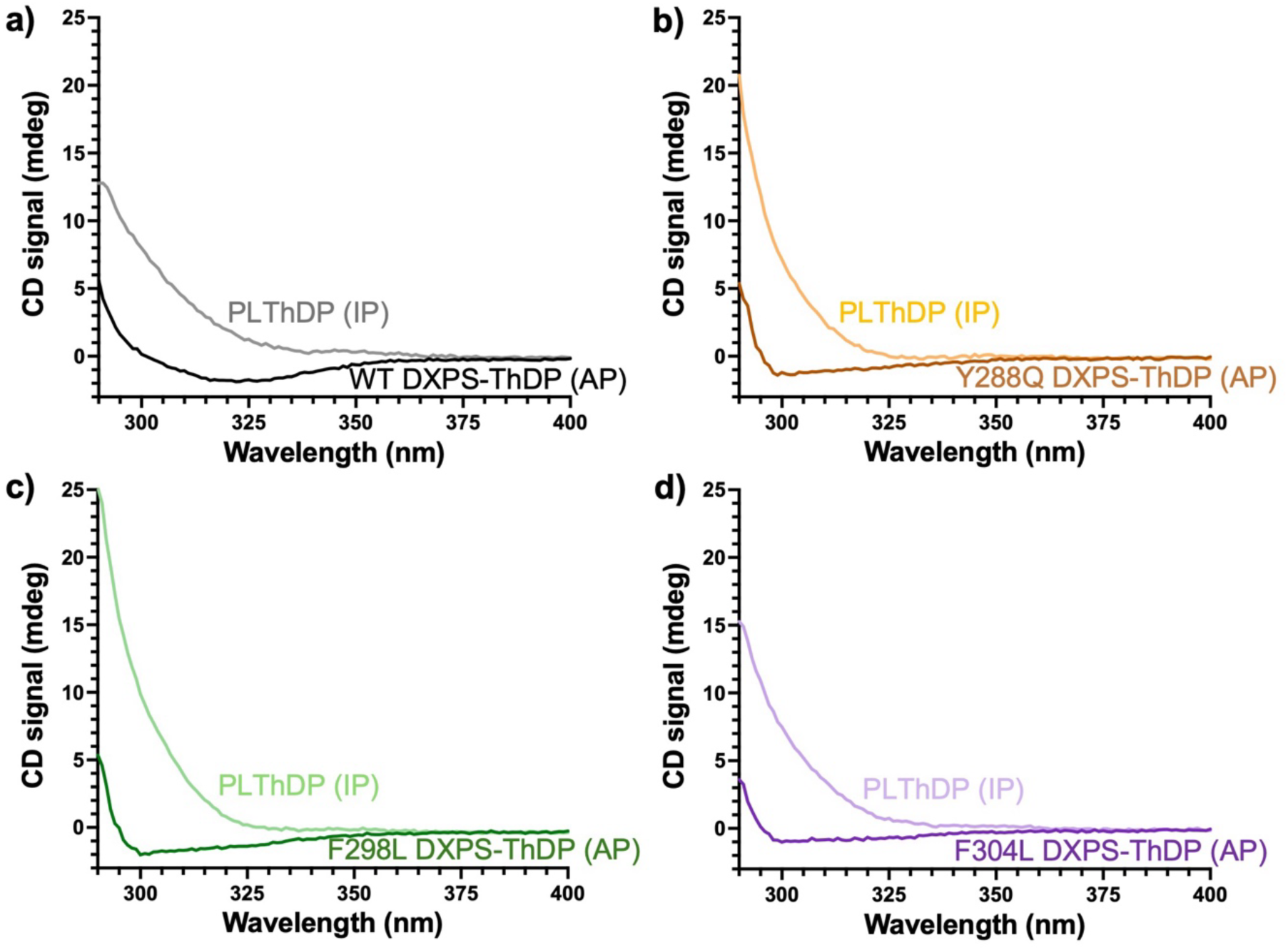
AP and PLThDP signals on WT and DXPS variants. CD scan of WT DXPS (a), Y288Q (b), F298L (c), and F304L (d) bound to ThDP or PLThDP, formed from the addition of 5 mM MAP.

**Figure S7:**
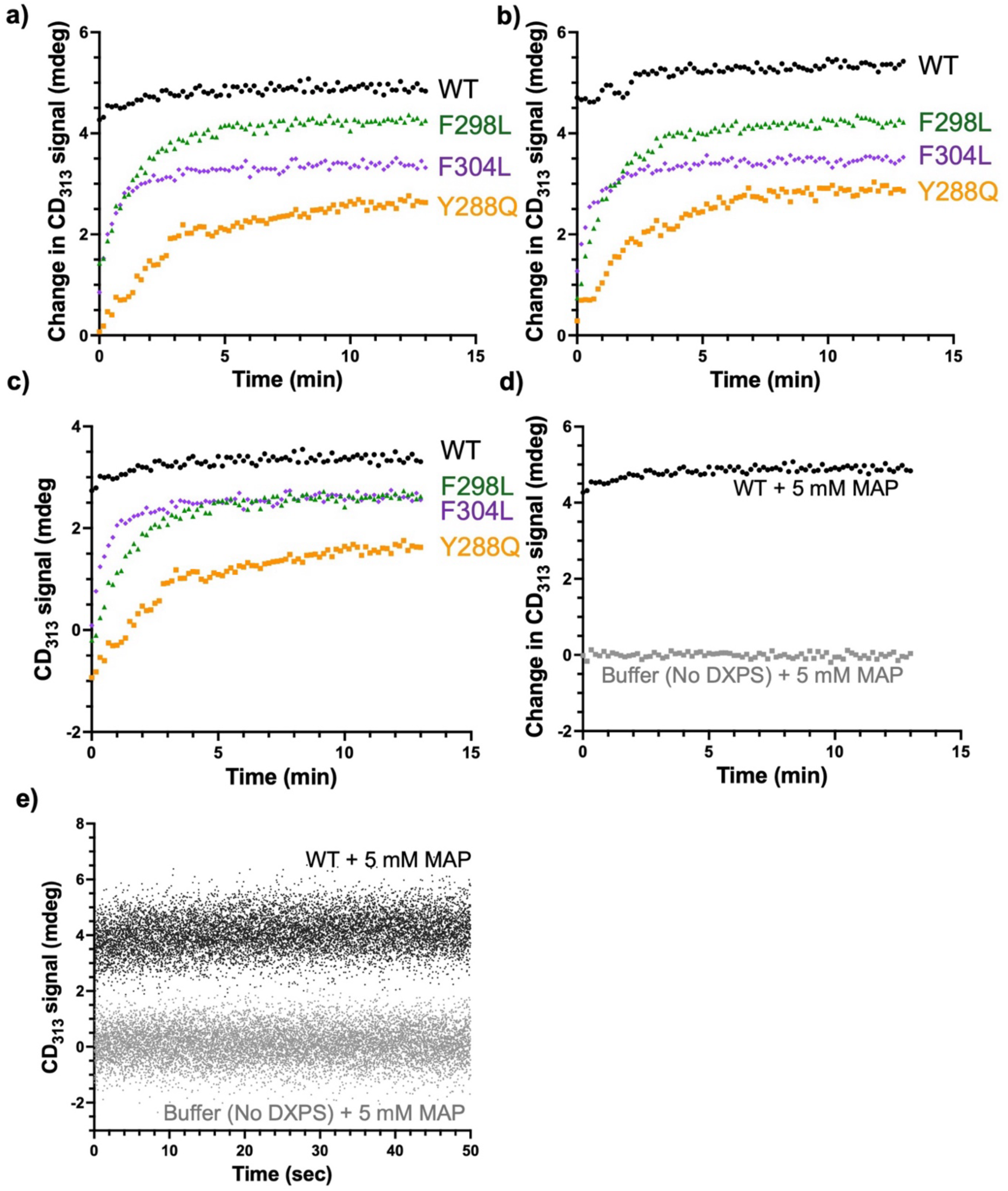
PLThDP formation on WT and DXPS variants. Accumulation of CD_313_ signal from AP signal subtracted, as shown in Figure 5d, (a) and duplicate data (b) observing PLThDP formation by monitoring the change in CD_313_ signal over 13 minutes. c) Unnormalized data from Figures 5d and S7a monitoring PLThDP signal accumulation. (d,e) Comparison of the PLThDP signal observed when 5 mM MAP is added to WT DXPS (black) and or buffer (grey, no DXPS control) in time resolved (d) and normalized, steady-state (e) CD experiments.

**Figure S8:**
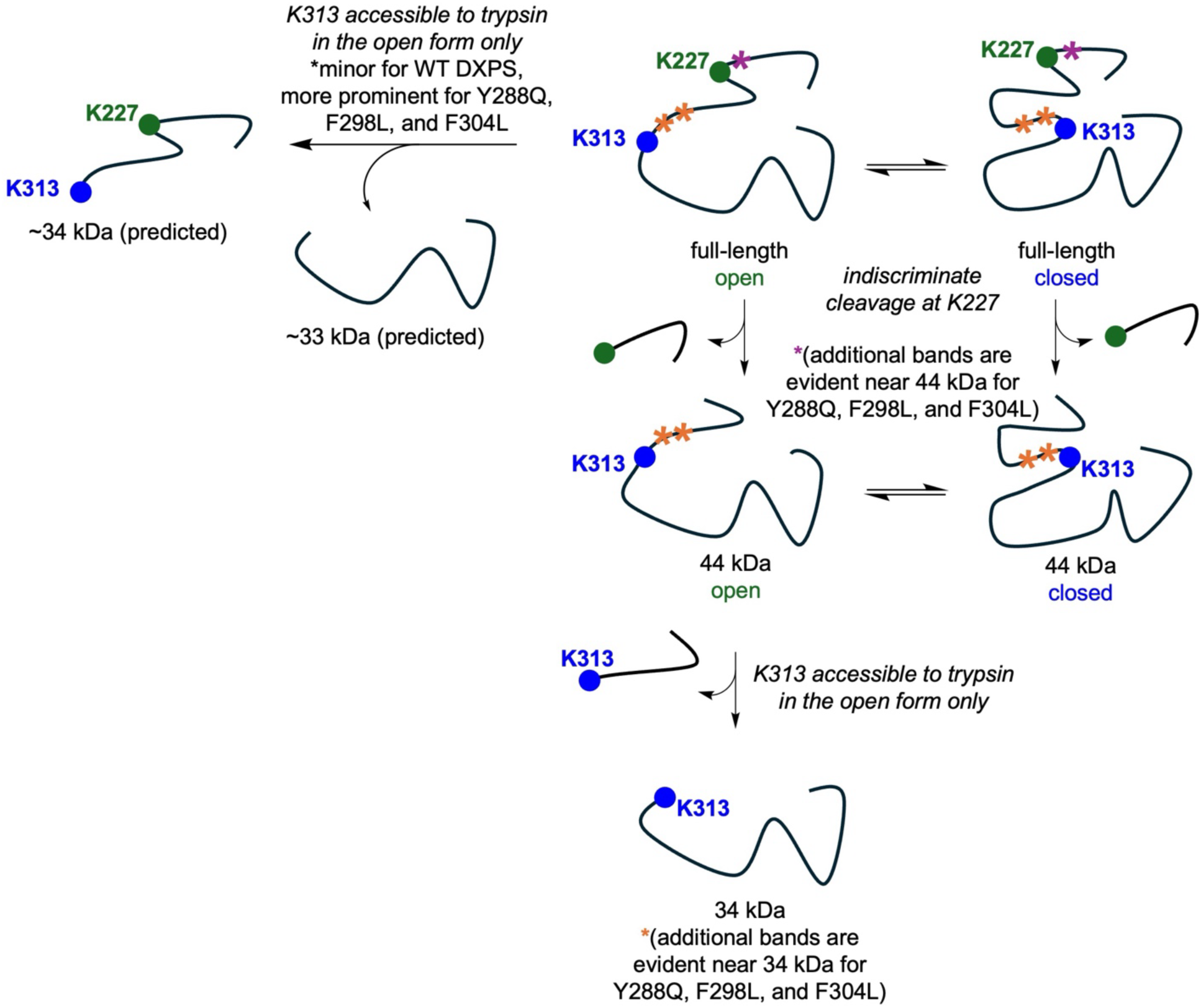
Trypsinolysis captures shifts in conformational equilibria of WT and variant *Ec*DXPS. Two major cleavage products were previously identified on the WT DXPS with accessibility dependent on global enzyme conformation.^2^ Cleavage of K227 occurs indiscriminately on either conformation, producing a 44 kDa peptide fragment. However, cleavage of K313 in the fork motif occurs only upon formation of the open conformation where the fork motif is disordered. Y288Q, F298L, and F304L appeared to exist predominately in an open conformation, evidenced by the rapid accumulation of the 34 kDa peptide. Additionally, digestion of each variant led to “doublet” bands near the 34 and 44 kDa major cleavage sites. We reason that a predominately open enzyme may be cleaved at K313 before or after K227, which would produce 34 and 33 kDa peptides. It is also plausible that more open variant enzymes may be more susceptible to cleavage a nearby Arg and Lys residues, resulting in the observed doublet bands. For example, cleavage at K203 is predicted to produce a 46 kDa peptide (*) while K286 or K273 digestion would produce 36 and 38 kDa bands, respectfully (*).

**Figure S9:**
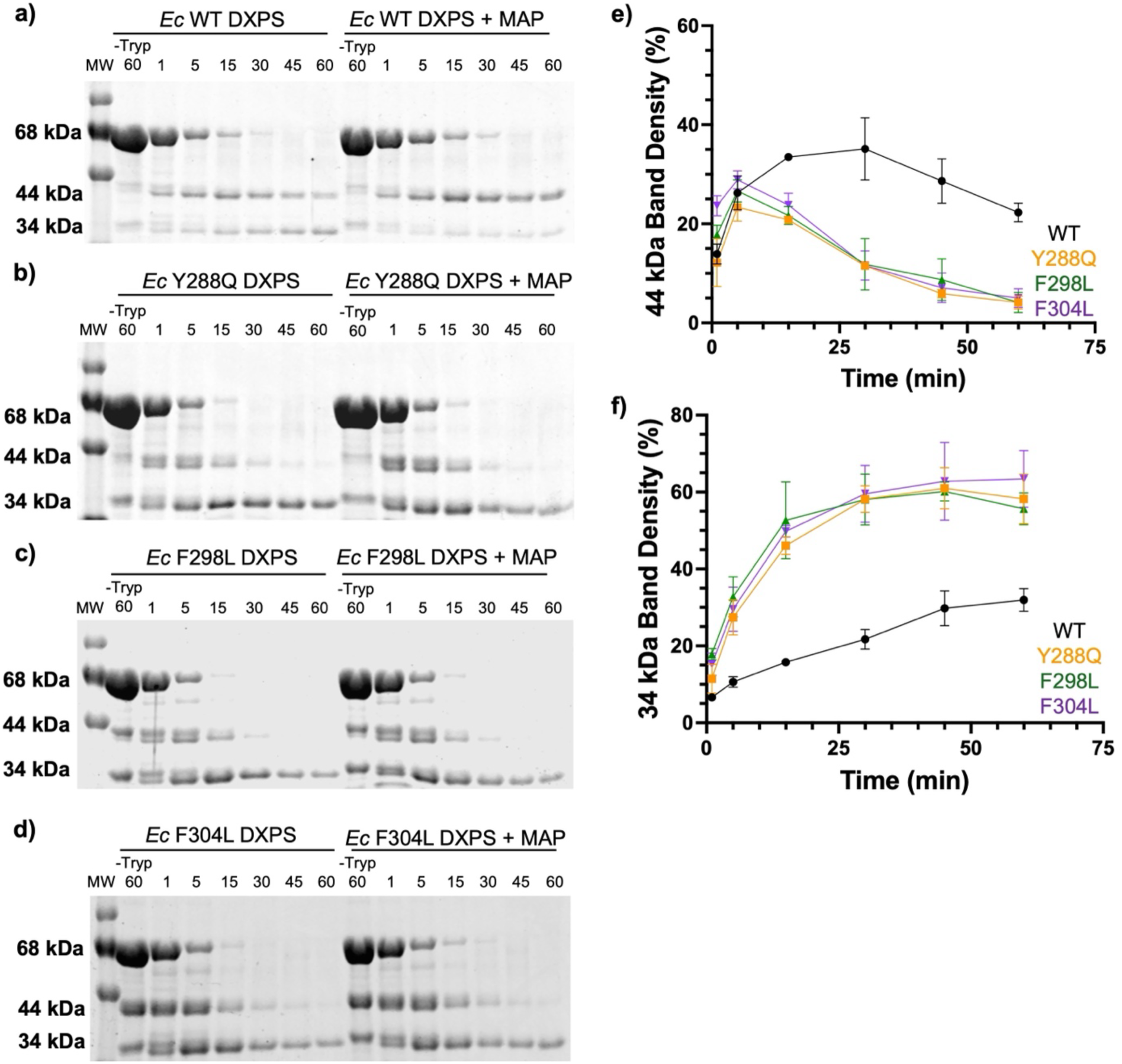
Limited trypsinolysis on WT and variant DXPS in the presence or absence of MAP. Representative trypsin digests are shown for (a) WT (as presented in Figure 7a), (b) Y288Q (shown in Figure 7b), (c) F298L, and (d) F304L cleavage products over 60 minutes, in the absence (left) or presence (right) of 5 mM MAP; e) Densitometry analysis of the 44 kDa and minor second band indicates this peptide accumulates to lower levels and is more rapidly converted to 33/34 kDa peptides on all DXPS variants in the absence of ligand; f) Quantification of the 34 kDa band shows rapid accumulation of all variants, relative to WT DXPS in the absence of MAP.

**Figure S10:**
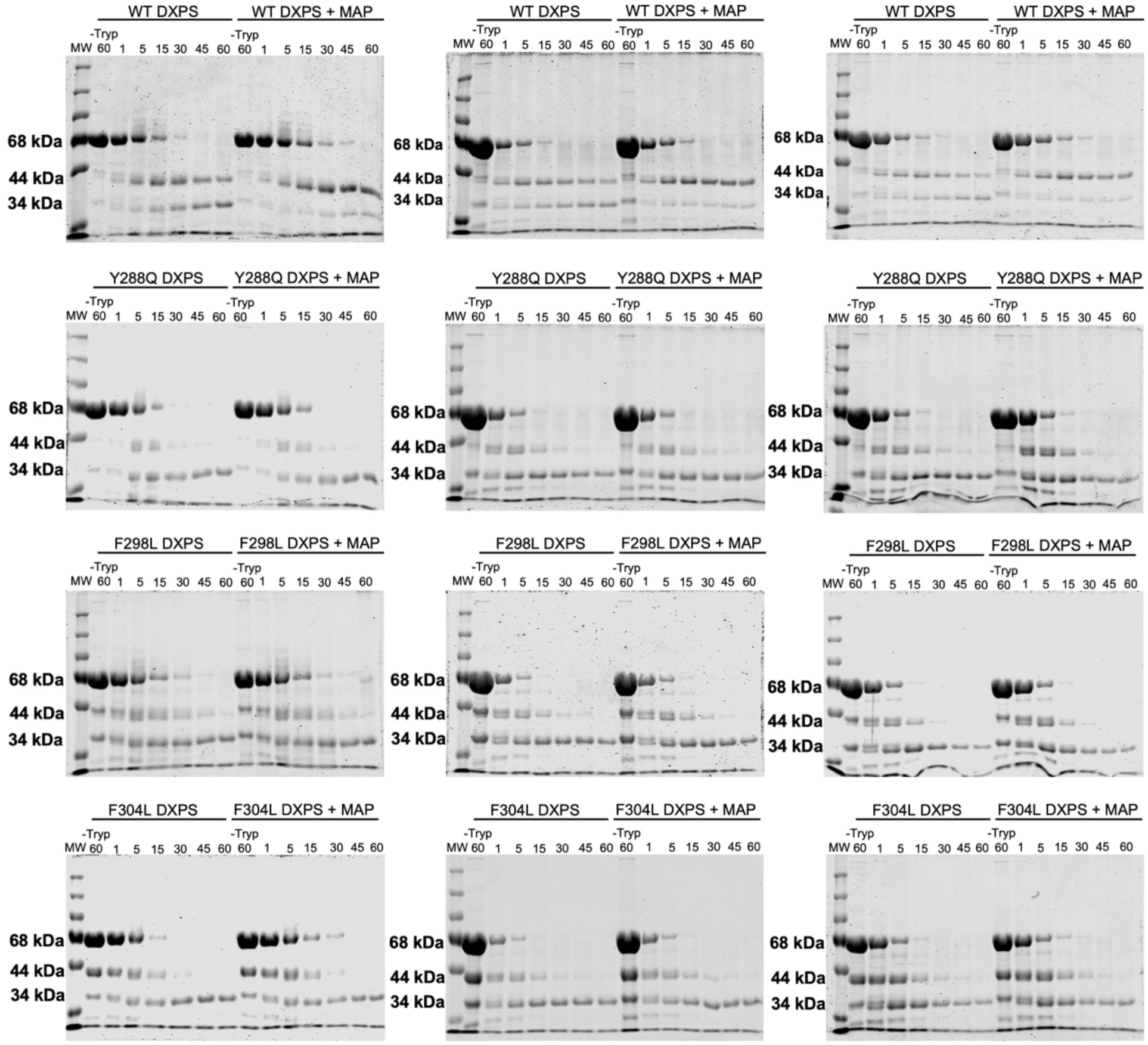
Gels showing replicate limited trypsinolysis experiments. Time courses of raw, unedited trypsin digests are shown for WT, Y288Q, F298L, and F304L over 60 minutes, as shown in Figures 6 and S9.

**Figure S11:**
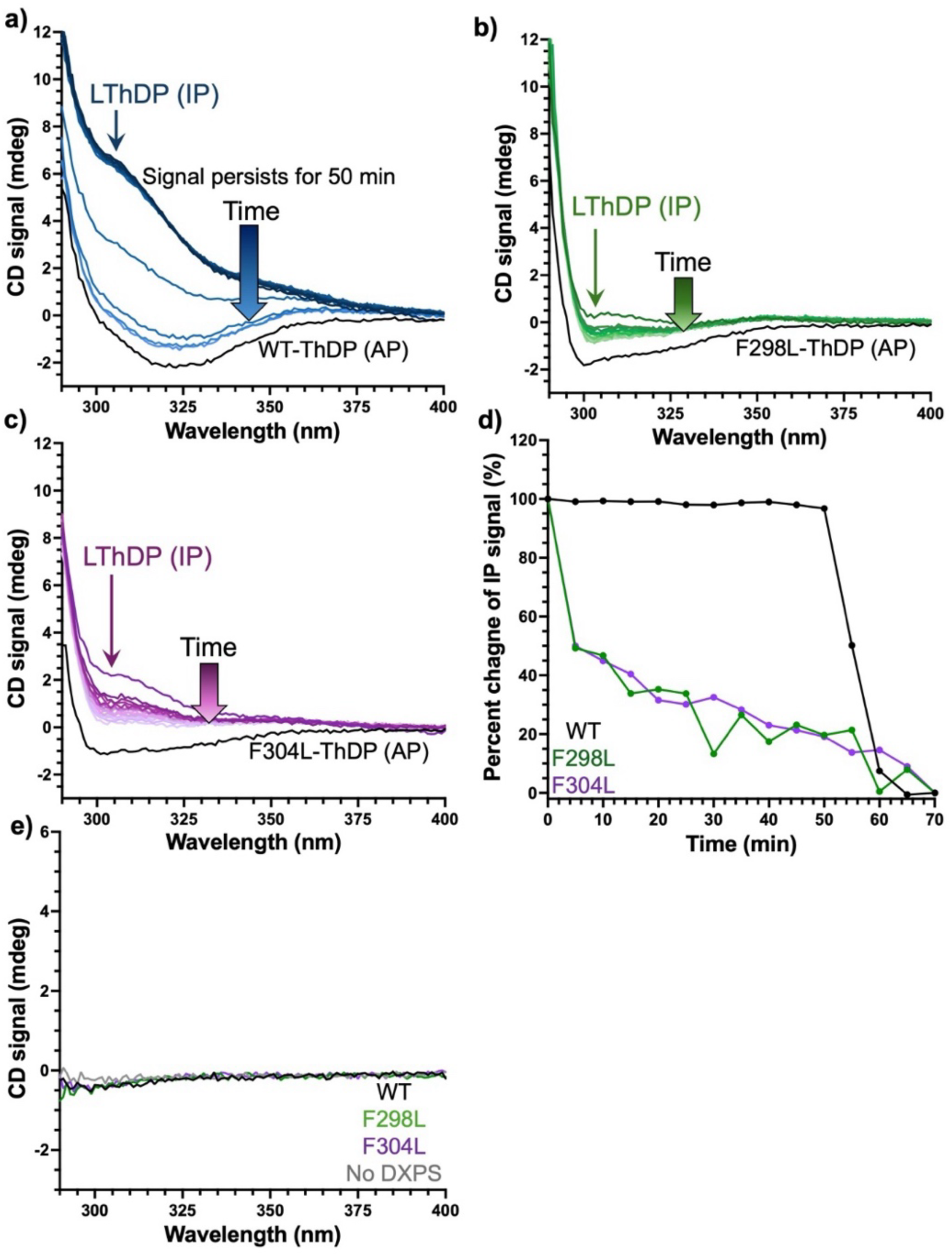
Duplicate CD data showing LThDP persistence (anaerobic) on (a) WT, (b) F298L, and (c) F304L DXPS. LThDP was formed upon addition of 5x*K*_m_^Pyr^. Intermediate was measured for 70 min where a new equilibrium observed upon signal depletion. The AP signal of each enzyme is shown in black. (d) Comparing percent change in IP signal depletion on WT, F298L, and F304L DXPS enzymes. (e) Enzyme was removed by filtration, and the clarified solution was scanned 290-400 nm. Representative replicates in which a chiral product was not detected from pyruvate decarboxylation on WT or DXPS variants are shown. Upon depletion of the LThDP IP signal in persistence experiments, the reaction mix was returned to the anaerobic chamber where it was added to 3 kDa microfuge filters. Enzymes were filter separated by centrifugation at 14,000 x g at room temperature, and the remaining solution was added back to the 1.5 mL, 10 mm path-length cuvette (Starna cells, 1-Q-10-ST-S). A steady state CD scan was recorded of each filtrate at 25 °C from 290-400 nm with a 1 nm step and 0.5 s averaging time to determine the presence of a chiral product.

**Figure S12:**
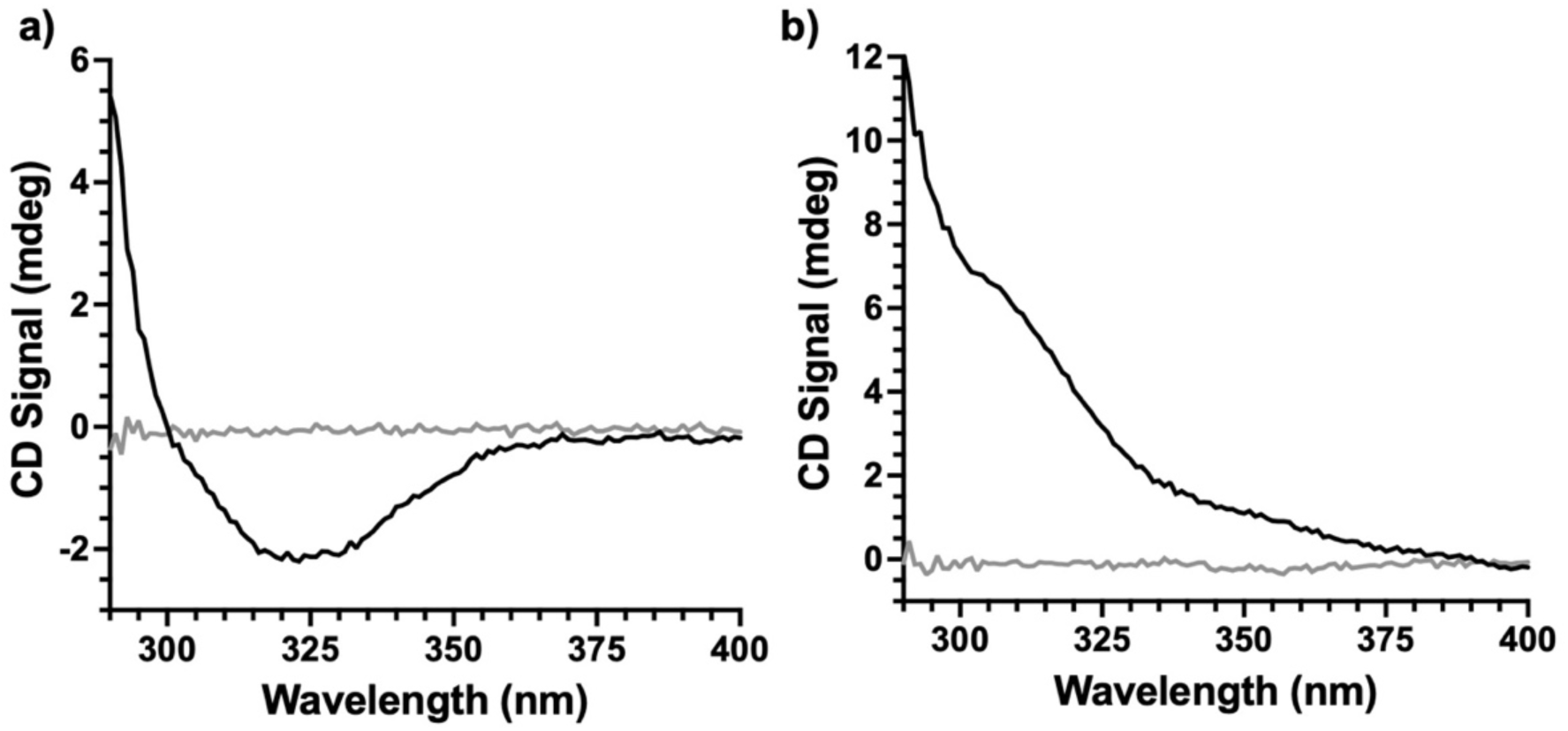
(a) Representative replicates comparing WT DXPS (black) and dialysis buffer (grey) alone, and (b) with 5*K*_m_^Pyr^ to WT and 5mM added to the dialysis buffer sample to observe the AP and IP signatures respectfully. Refer to Figure 5a,b which first showed WT AP and IP signals.

**Figure S13:**
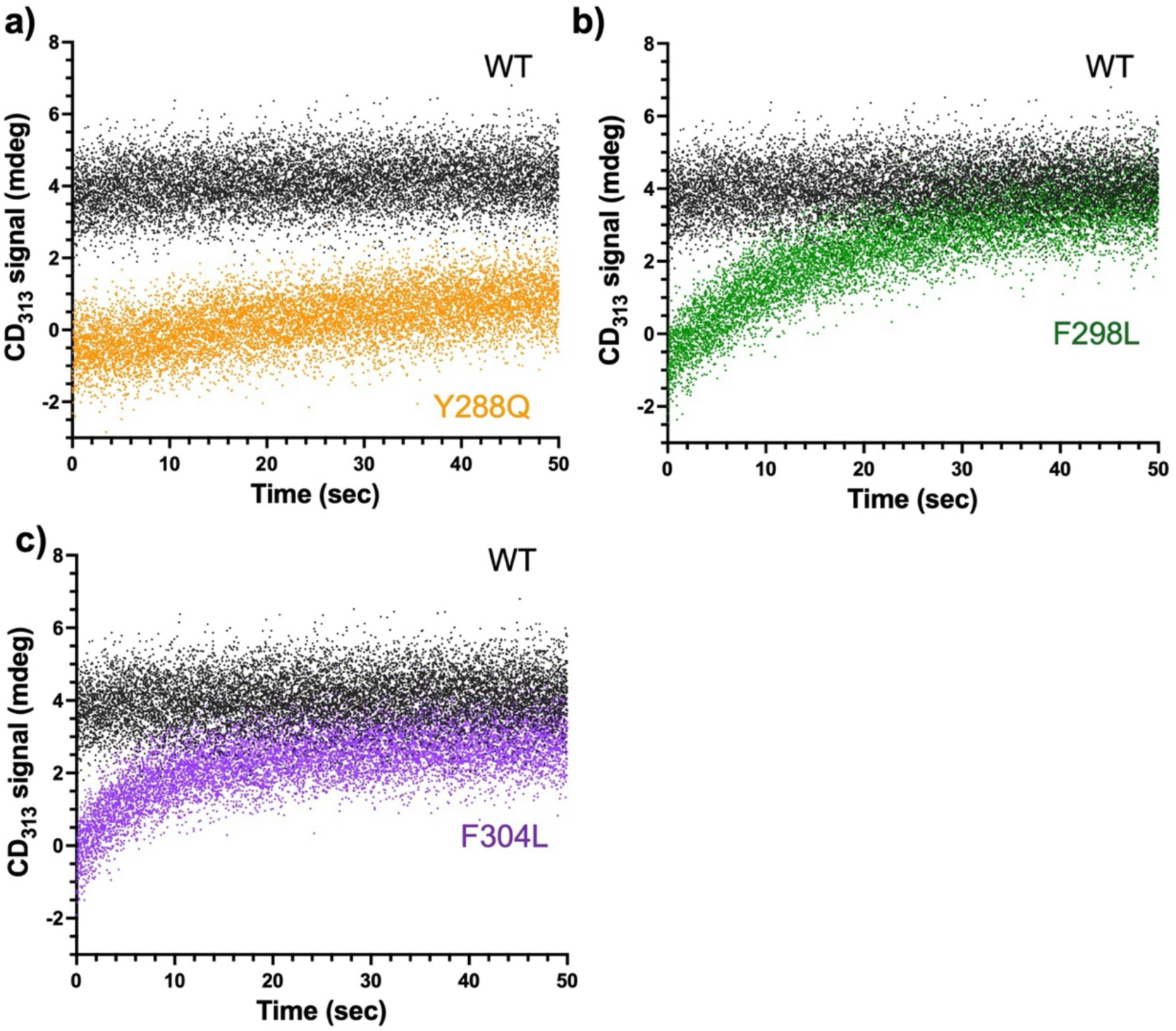
Disruption of aromatic cluster leads to slower PLThDP formation at 25 °C. Time-resolved experiments monitoring of the PLThDP CD_313_ signal upon addition of MAP at 25 °C revealed slower PLThDP formation when the aromatic cluster is disrupted on (a) Y288Q (orange), (b) F298L (green), and (c) F304L (purple) compared to WT DXPS (black) under conditions similar to those used to detect HCO_3_^−^ and HEThDP by NMR. The slow intermediate formation observed at 10 °C necessitated preparation of NMR samples at ambient temperatures. Thus, we followed similar methods to compare PLThDP formation in complementary conditions to the NMR sample preparation which was used to observe the fate of pyruvate. Briefly, 60 µM WT or variant DXPS was prepared in reaction buffer containing 50 mM HEPES, pH 8, 100 mM NaCl, 2 mM MgCl_2_, and 2 mM ThDP. A 10 mM MAP solution was prepared in a separate reaction buffer. Using the CD attached to the stopped-flow SF3 accessory, enzyme and MAP solutions (30 µM enzyme and 5 mM MAP final concentrations) were rapidly mixed. The CD_313_ signal was monitored over 50 s at 25 °C, and the data from 7 repetitive shots were averaged.

**Figure S14:**
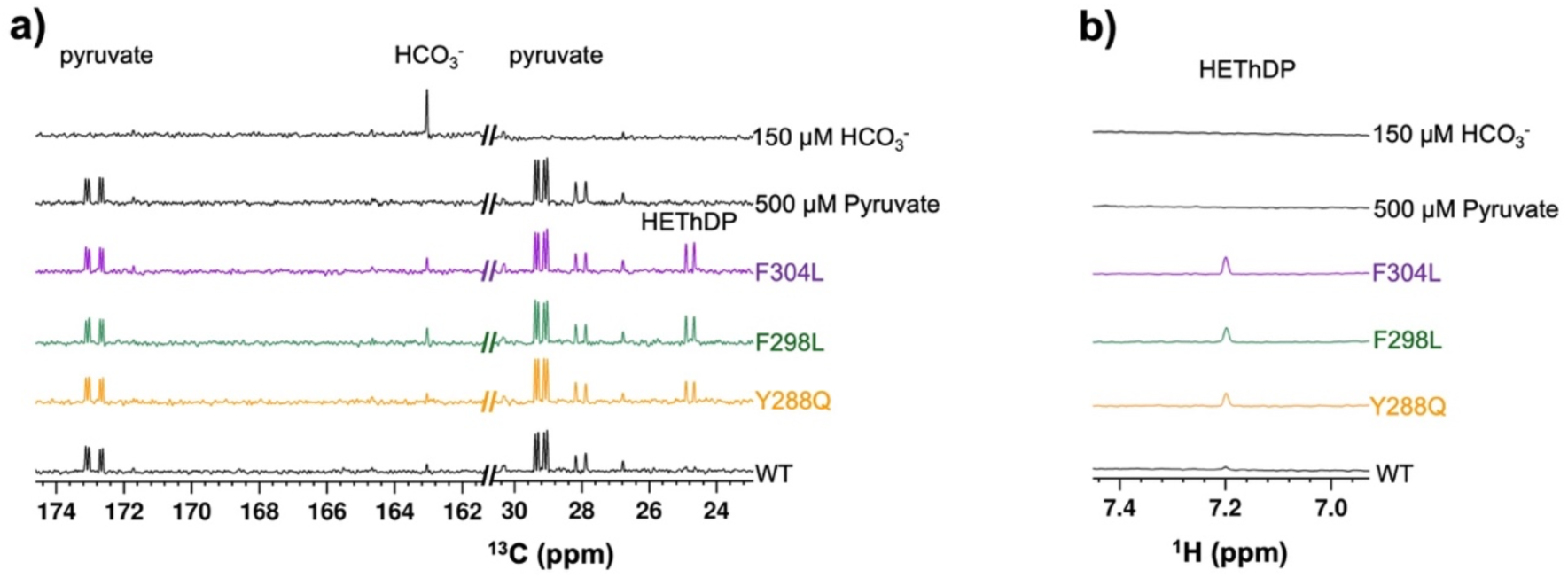
Observation of HEThDP ejection from Y288Q, F298L, and F304L DXPS. Duplicate data are shown of ^13^C (a) and ^1^H (b) spectra showing HCO_3_^−^ and HEThDP formation upon disruption of the aromatic cluster compared to the pyruvate and HCO_3_^−^ standards.

**Figure S15:**
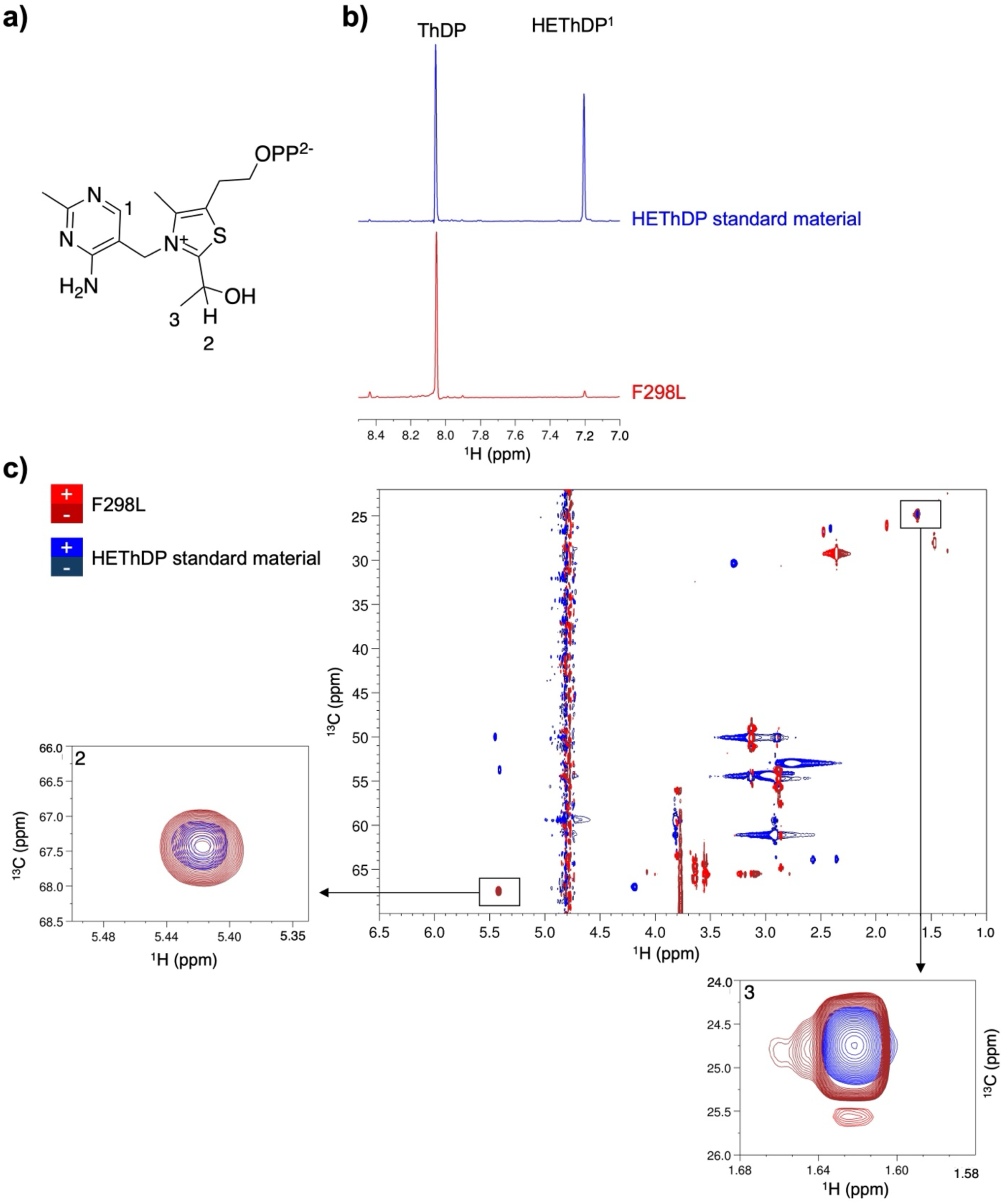
HEThDP identification in the DXPS variant representative F298L samples. (a) Structure of HEThDP; (b) ^1^H NMR spectra comparing the presence of a 6’-AP ring proton at ∼7.2 ppm (1) between the standard material (blue) and the unknown in the F298L (red) sample;^4,5^ (c) Using the natural abundance of ^13^C in the 1.3 mg/mL HEThDP standard material, a two-dimensional HSQC experiment revealed overlaying components of the hydroxyethyl group corresponding to –C_2ɑ_-H (2) and -C_2ɑ_-CH_3_ (3) species at (5.42, 67.5) and (1.62, 24.77) ppm, respectively, between the standard and the F298L unknown sample. Together these data support HEThDP production in the F298L sample. Peaks of interest are numerically labeled on the HEThDP structure in panel a.

Two-dimensional NMR samples were prepared as described in the materials and methods for the detection of bicarbonate and HEThDP by one dimensional NMR. A HEThDP standard solution was prepared in the NMR reaction buffer at a final concentration of 1.3 mg/mL.^3^ The natural abundance of ^13^C was used to compare to the unknown product in the F298L sample chosen as a representative from the aromatic disrupting variant series. Two-dimensional NMR data was collected on a Bruker Avance spectrometer fitted with a TCI cyrogenic probe including a cyro-cooled ^13^C preamplifier.

The HEThDP ^1^H-^13^C HSQC was acquired at natural abundance of ^13^C using a gradient based experiment with the following parameters:

| Parameter | $^1\text{H}$ | $^{13}\text{C}$ |
| --- | --- | --- |
| Carrier (ppm) | 4.77 ($\text{H}_2\text{O}$ ppm @ 25 °C) | 46.24 |
| Spectral Width (ppm) | 16 | 50 |
| Acquisition time/FID (ms) | 125 | 15 |
| Complex data points | 1200 | 114 |

Eight scans/FID and a recycle delay of 1.3 s were used, for a total data collection time of 45 minutes.

The F298L ^1^H-^13^C correlated spectrum is a modified gradient-HSQC experiment designed to (1) select for CH and CH_3_ groups and eliminate CH_2_ signals and (2) acquired in constant-time mode along ^13^C so that ^1^H-^13^C-^13^C signals have opposite phase relative to ^1^H-^13^C-^12^C peaks. Data collection in this mode was used to confirm the presence of the hydroxyethyl ^13^CH_3_-^13^C_2a_H-OH moiety. Data acquisition parameters are as follows:

| Parameter | $^1\text{H}$ | $^{13}\text{C}$ |
| --- | --- | --- |
| Carrier (ppm) | 4.77 ($\text{H}_2\text{O}$ ppm @ 25 °C) | 46.24 |
| Spectral Width (ppm) | 14 | 50 |
| Acquisition time/FID (ms) | 125 | 15 |
| Complex data points | 1077 | 114 |

Sixteen scans/FID and a recycle delay of 1.3 s were used, for a total data collection time of 1.5 hrs.

**Figure S16:**
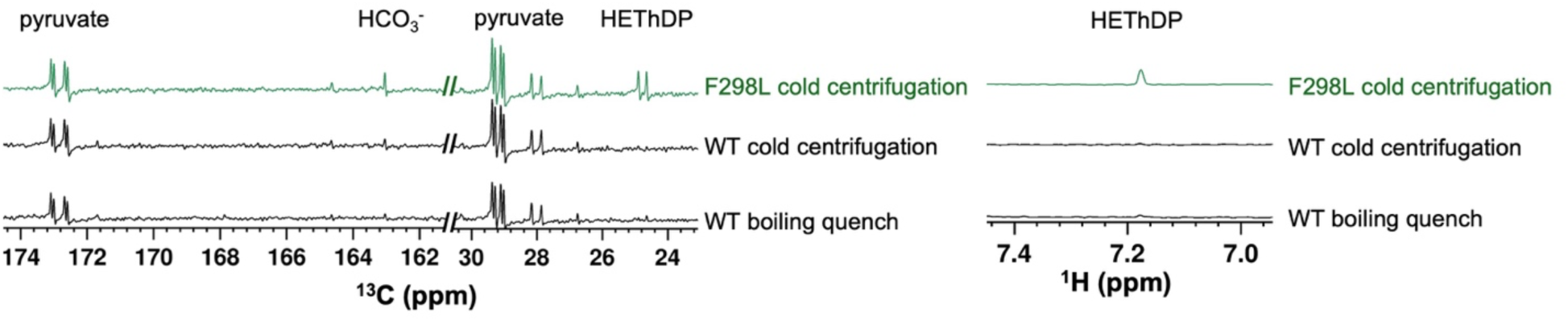
Discernment between pre- and post-decarboxylation intermediate by NMR ^13^C (left) and ^1^H (right) spectra showing accumulation of intermediate in solution in the presence of F298L DXPS when quenching by the 4 °C centrifugation method to remove enzyme using a 10 kDda filter from the sample solution. Samples were prepared in a similar manner to those described in the materials and methods to detect bicarbonate and HEThDP, with a change in the quenching method. Briefly, solutions were degassed in the anaerobic chamber, and WT or variant DXPS (5 µM) was added to NMR reaction buffer (50 mM HEPES, pH 8; 100 mM NaCl; 2 mM MgCl_2_; 1 mM ThDP; and 5 % D_2_O). Addition of 500 µM [^13^C_3_]pyruvate to a final reaction volume of 700 µL initiated the formation of LThDP. Samples were incubated at ambient chamber temperature (∼27 °C) for 20 min. A control WT DXPS sample was quenched by boiling at 95 °C for 5 min prior to filtration using a 10 kDa microfuge filter tube by centrifugation at 14,000 x g for 10 min. A F298L DXPS sample in a filter microfuge tube was sealed with parafilm, removed from the chamber, and immediately placed on ice. The sample was centrifuged at 16,000 x g for 25 min at 4 °C in a cold room. A WT DXPS control sample was prepared as described for F298L. The filtrates in each case (600 µL) were added to prechilled microfuge tubes containing 15 µL of 10 mM gadobutrol (MedChem Express, NJ) to enhance the ^13^C signal of HCO_3_^−^. The solutions were then transferred to chilled NMR tubes and stored at 4 °C until NMR acquisition which was performed the same day as sample preparation. Spectra were acquired at 150 MHz (^13^C), at 25 °C, using a 35^°^ ^13^C excitation pulse, 1,320 scans/FID, 150 ms acquisition time/FID, and a 3s relaxation between scans.

**Figure S17:**
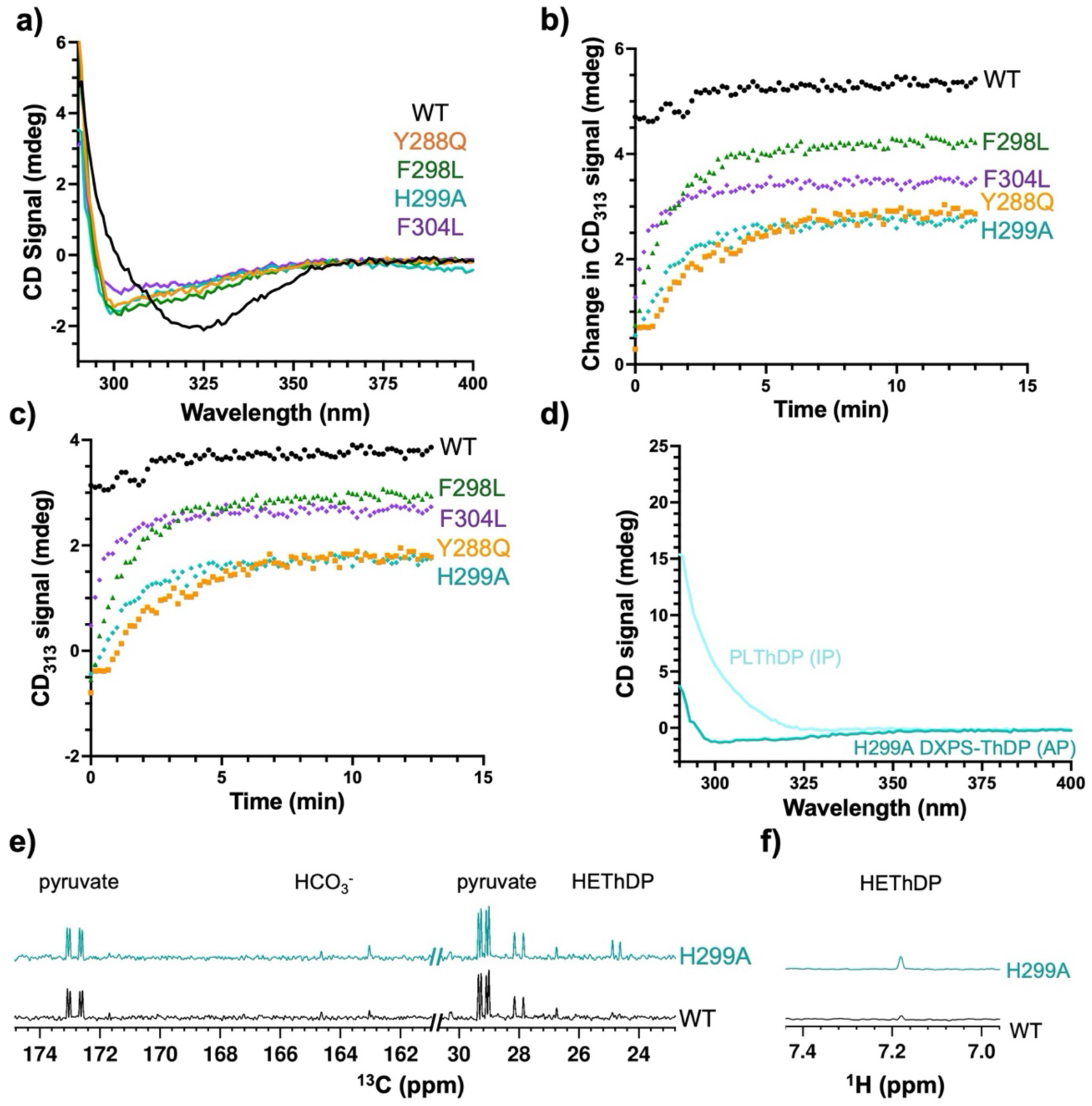
Duplicate data of H299A PLThDP formation overlaid with aromatic disrupting variants depicting H299A AP signal (a), normalized (b) and unnormalized (c) aerobic PLThDP formation, as shown first in Figure S7, comparing WT and aromatic disrupting variant, observation of the H299A AP and PLThDP IP signals (d), ^13^C spectra (e), and ^1^H spectra (f).

